# Lung structural cells are altered by influenza virus leading to rapid immune protection following re-challenge

**DOI:** 10.1101/2024.07.20.604410

**Authors:** Julie C. Worrell, Kerrie E. Hargrave, George E. Finney, Chris Hansell, John Cole, Jagtar Singh Nijjar, Fraser Morton, Marieke Pingen, Tom Purnell, Kathleen Mitchelson, Euan Brennan, Jay Allan, Georgios Ilia, Vanessa Herder, Claire Kennedy Dietrich, Yoana Doncheva, Nigel B. Jamieson, Massimo Palmarini, Megan K.L MacLeod

## Abstract

Lung structural cells, including epithelial cells and fibroblasts, form barriers against pathogens and trigger immune responses following infections such as influenza A virus. This response leads to the recruitment of innate and adaptive immune cells required for viral clearance. Some of these recruited cells remain within the lung following infection and contribute to enhanced viral control following subsequent infections. There is growing evidence that structural cells can also display long-term changes following infection or insults. Here we investigate long-term changes to mouse lung epithelial cells, fibroblasts, and endothelial cells following influenza virus infection and find that all three cell types maintain an imprint of the infection, particularly in genes associated with communication with T cells. Lung epithelial cells from IAV-infected mice display functional changes by more rapidly controlling influenza virus than cells from naïve animals. This rapid anti-viral response and increased expression of molecules required to communicate with T cells demonstrates sustained and enhanced functions following infection. These data suggest lung structural cells could be effective targets for vaccines to boost durable protective immunity.

**Graphical Abstract:** 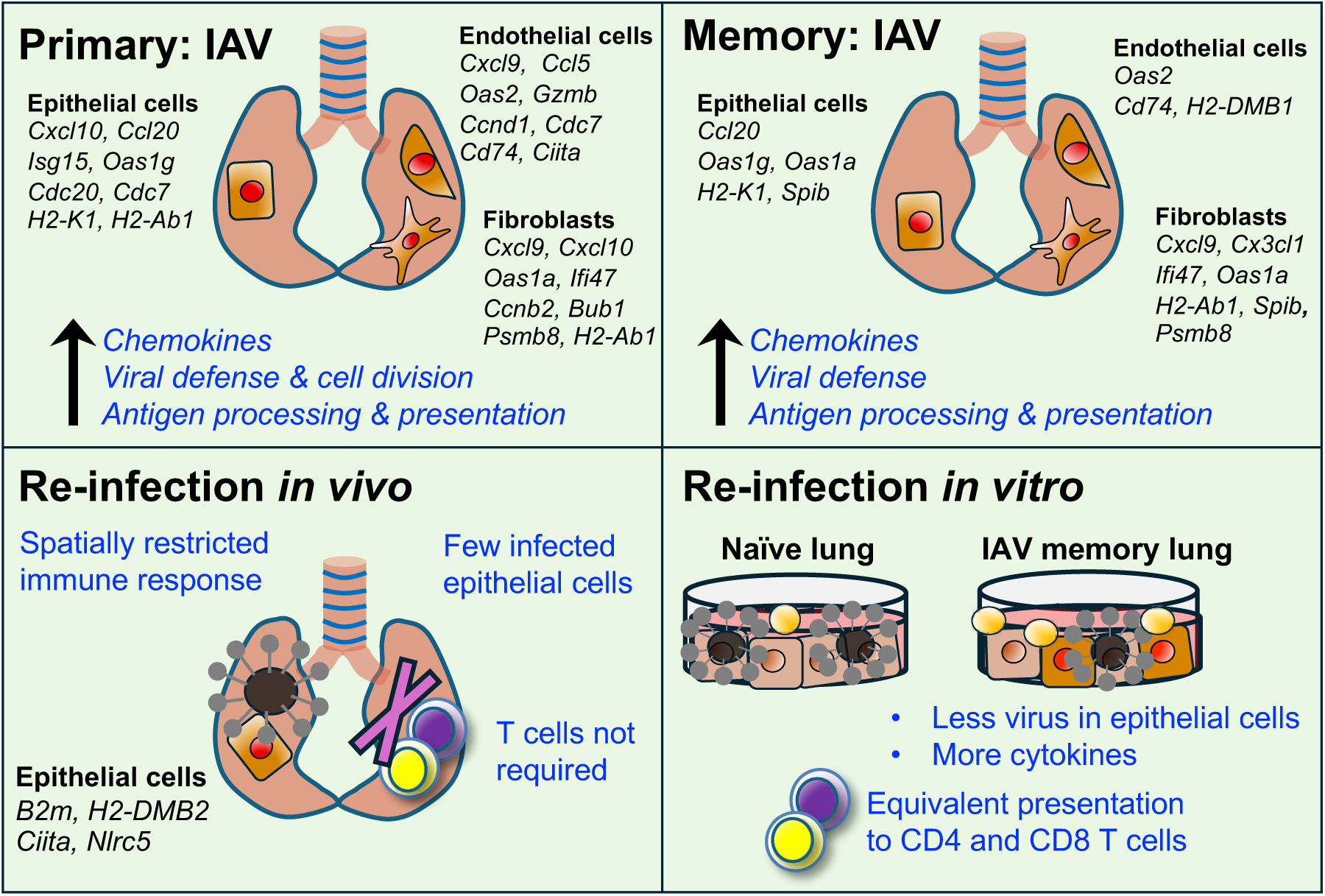

**Highlights:**

- Lung epithelial cells, fibroblasts, and blood endothelial cells maintain an inflammatory imprint of influenza A virus (IAV) infection for at least 40 days post-infection.
- *In vivo* re-infection leads to a more spatially restricted anti-viral response compared to primary IAV-infected animals.
- T cells are not required for enhanced viral control early after re-infection *in vivo*
- *Ex vivo* lung epithelial cells from IAV-infected mice more rapidly control IAV than cells from naïve animals in the absence of immune cells.

## Introduction

The lung is constantly exposed to microbes that can trigger immune responses. Respiratory viruses, including influenza A virus (IAV), are leading causes of disease worldwide^1,2^. The World Health Organization estimates annual epidemics result in approximately 3 to 5 million cases of severe illness and about 290,000 to 650,000 deaths.

Many different cell types are required to control the pathogens that enter the lung. These include structural cells, epithelial cells and stromal cells (fibroblast and endothelial cells); and immune cells that can be resident or recruited following the infection. The early responses to IAV by lung structural cells have been well characterized^3–8^. Epithelial cells are the primary cell type infected *in vivo* and are the cells most likely to produce new virus^9,10^. They, and nearby fibroblasts, trigger an early wave of inflammatory cytokines and chemokines that can help control the infection until dedicated immune cells migrate to the infection site^3–6^. Blood endothelial cells (BECs) play a key role in alerting circulating immune cells and recruiting them to the lung via chemokines^7,11^.

IAV can be cleared within 7 to 10 days, but the consequences of the infection are more sustained. While epithelial cells and fibroblasts are involved in tissue repair, damage can persist for many months in mouse infection models ^12–14^. Lung immune cells are also altered by infection: adaptive T and B cells can take up long-term residence in tissues as Tissue resident memory (Trm) cells and circulating monocytes can differentiate into alveolar macrophages. These cells can provide both pathogen specific and non-specific protection to future challenges and are considered arms of adaptive and innate memory^15–22^.

Recently, evidence is growing that structural cells can also display long-term changes to infections^23,24^. In the context of IAV, surfactant producing club epithelial cells that survive IAV infection can produce a rapid anti-viral response following a second challenge which may contribute to tissue damage^25,26^. Other epithelial cells can also survive IAV infection by reducing communication with lung CD8 T cells^27^. However, an in-depth analysis of long-term changes by IAV to lung epithelial cells is lacking. Moreover, little is known about the long-term changes to other lung structural cells following influenza virus infection.

To address these questions, we used transcriptomic and flow cytometry analysis to investigate long-term changes to lung structural cells following primary infection and re-challenge of mice with IAV. Our data reveal that, in addition to epithelial cells, fibroblasts and BECs maintain an imprint of the IAV infection. Our analysis demonstrates that antigen processing and presentation is a shared pathway upregulated in all three cell types for at least 40 days following infection and implicates the Ets family transcription factor, SpiB^28^, in regulating these changes.

More rapid control of IAV is expected in re-challenge infections, even when different IAV strains are used in a heterosubtypic infection to avoid neutralizing antibodies^18,21,29,30^. We find that lung epithelial cells rapidly upregulate the expression of molecules required for communication with T cells and present antigen to IAV-specific CD4 and CD8 T cells *in vitro*. Despite this ability to communicate with lung T cells, *in vivo* rapid virus control within the first two days of a re-infection is independent of T cells. Instead, our data show that epithelial cells from previously infected mice can more rapidly control IAV *in vitro* independently of immune cells. Together, our data suggest that lung structural cells maintain an imprint of IAV infection that can enhance their ability to communicate with T cells and for epithelial cells, also bolsters their own immunity against subsequent infections.

## Results

### IAV infection causes prolonged up-regulation of immune-related genes in lung structural cells

To advance our understanding of the long-term consequences of IAV infection on lung structural cells, we used fluorescence activated cell sorting (FACS) to isolate three major cell types in the lung: epithelium, fibroblasts, and endothelium. We sorted the cells based on the expression of EpCAM1 (epithelial cells), CD140a (fibroblasts), and CD31 (blood endothelial cells, BECs) from mice at day 10 (up to viral clearance) and 40 post-infection (long after viral clearance) and compared their gene expression to cells sorted from naïve animals. The gating strategy, sort purities and Principal Component Analysis (PCA) are shown in SFigA-E. PCA demonstrated substantial changes at day 10 post infection for all cell types compared to naïve controls. We identified 1016 differentially expressed genes (DEG) in epithelial cells (784 up, 232 down), 3128 DEG in fibroblasts (1772 up, 1356 down) and 1292 DEG in endothelial cells (968 up, 324 down) between day 10 IAV and naïve groups (Fig 1A-B and Extended data 1). Over representation analysis (ORA) detected enrichment in cell cycle in all three subsets and defense response to virus in epithelial cells and fibroblasts, SFig 2A-C.

**Figure 1:**
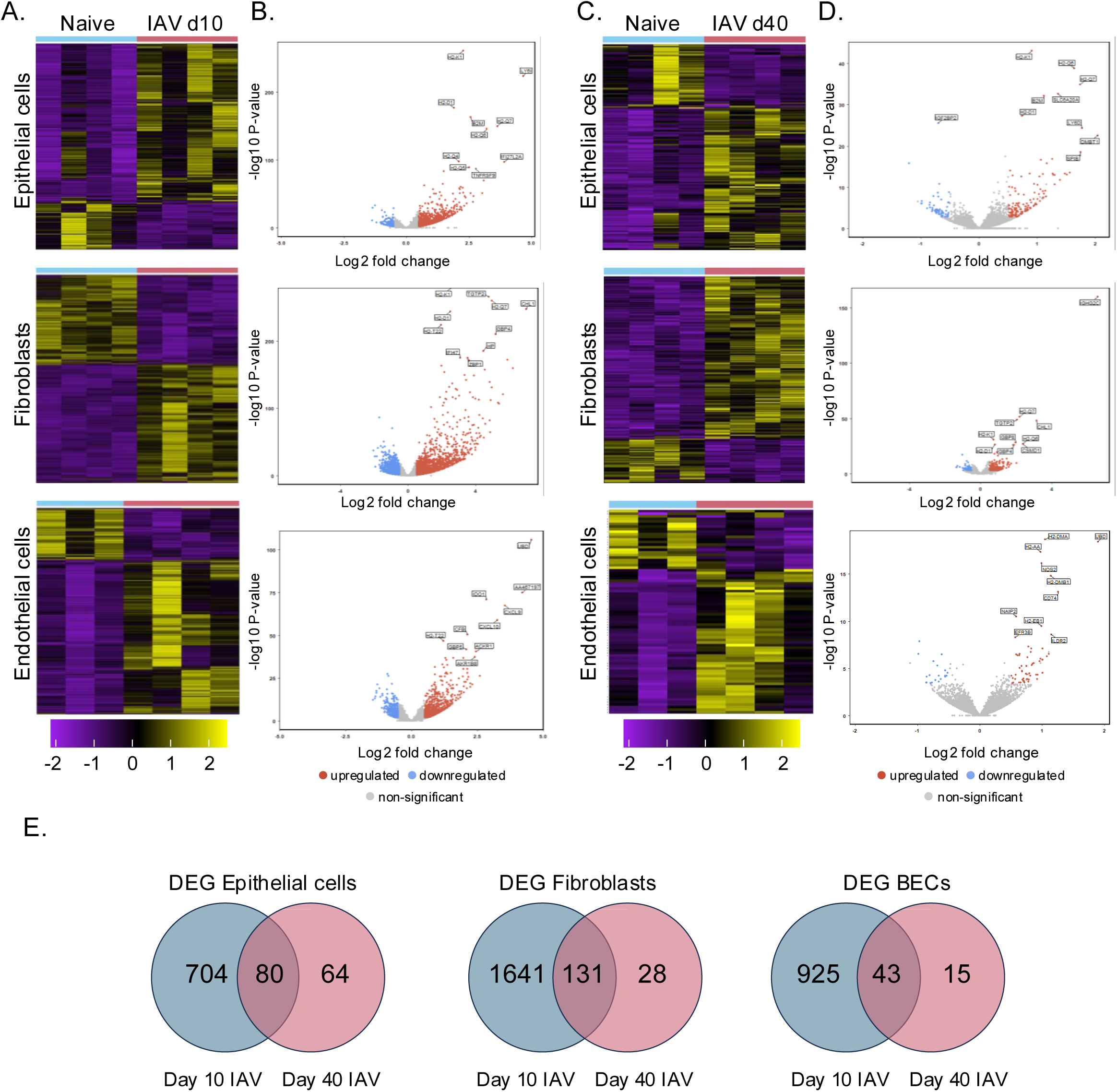
IAV infection induces conserved transcriptional changes at day 10 and 40 post infection. RNA sequencing was performed on sorted lung epithelial cells, fibroblasts, and endothelial cells from naïve and C57BL/6 mice infected with IAV 10 (A-B) or 40 (C-D) days previously. A, C: Differentially expressed genes (DEG) with a 2-fold change and an FDR q-value < 0.05 in lung structural cells were considered statistically significant. 7-8 mice were combined to make 4 samples per time point. B,D: Volcano plot showing the relationship between the fold change and the associated p−value with the top 10 DEG by fold change named. E: Venn diagrams showing overlap between differentially expressed upregulated genes at day 10 and 40 post IAV infection.

At day 40 post infection, all three cell types continued to show differential gene expression compared to cells from naïve mice, although the number of DEG were reduced compared to day 10: epithelial cells: 144 up, 61 down; fibroblasts 159 up, 43 down; BECs: 58 up and 21 down (Fig 1C-D). To validate the analysis, we isolated RNA from lung epithelial cells and fibroblasts in an independent experiment from naïve mice and animals infected 30 days earlier and compared gene expression in cells from naïve and infected animals by microarray (nCounter Nanostring mouse immunology panel with 29 custom genes, Extended data 2). We found a consistent fold change between DEG identified in the RNAseq that overlapped with genes in the microarray (SFig 3 and Extended data 2).

Closer inspection of overlapping upregulated DEG from our RNAseq datasets at day 10 and 40 post-infection, revealed that several genes remained persistently elevated with 144, 159 and 58 DEG in epithelial cells, fibroblasts, and BECs respectively (Fig 1E and Extended data 1). BECs had the lowest number of genes that remained upregulated, suggesting these cells may be more likely to return to homeostasis than epithelial cells or fibroblasts. ORA analysis of the DEG in the three cell types indicated sustained changes in pathways related to the innate and adaptive immune responses (SFig 2D-F). Taken together, these data reveal sustained transcriptional alterations in immune related genes in lung structural cells long after viral clearance.

We next performed a gene set enrichment network analysis on all structural cell types at both day 10 and 40 post infection (Fig 2A-C). This was performed as ORA can enrich for highly similar genes and may only represent a portion of the enriched biology. Network analysis revealed conserved upregulation of gene networks governing shared biological themes in all three cell types, including antigen processing and presentation at day 40 post IAV infection. These findings indicate that IAV infection leads to long-term changes in the expression of immune-related genes in lung structural cells.

**Figure 2:**
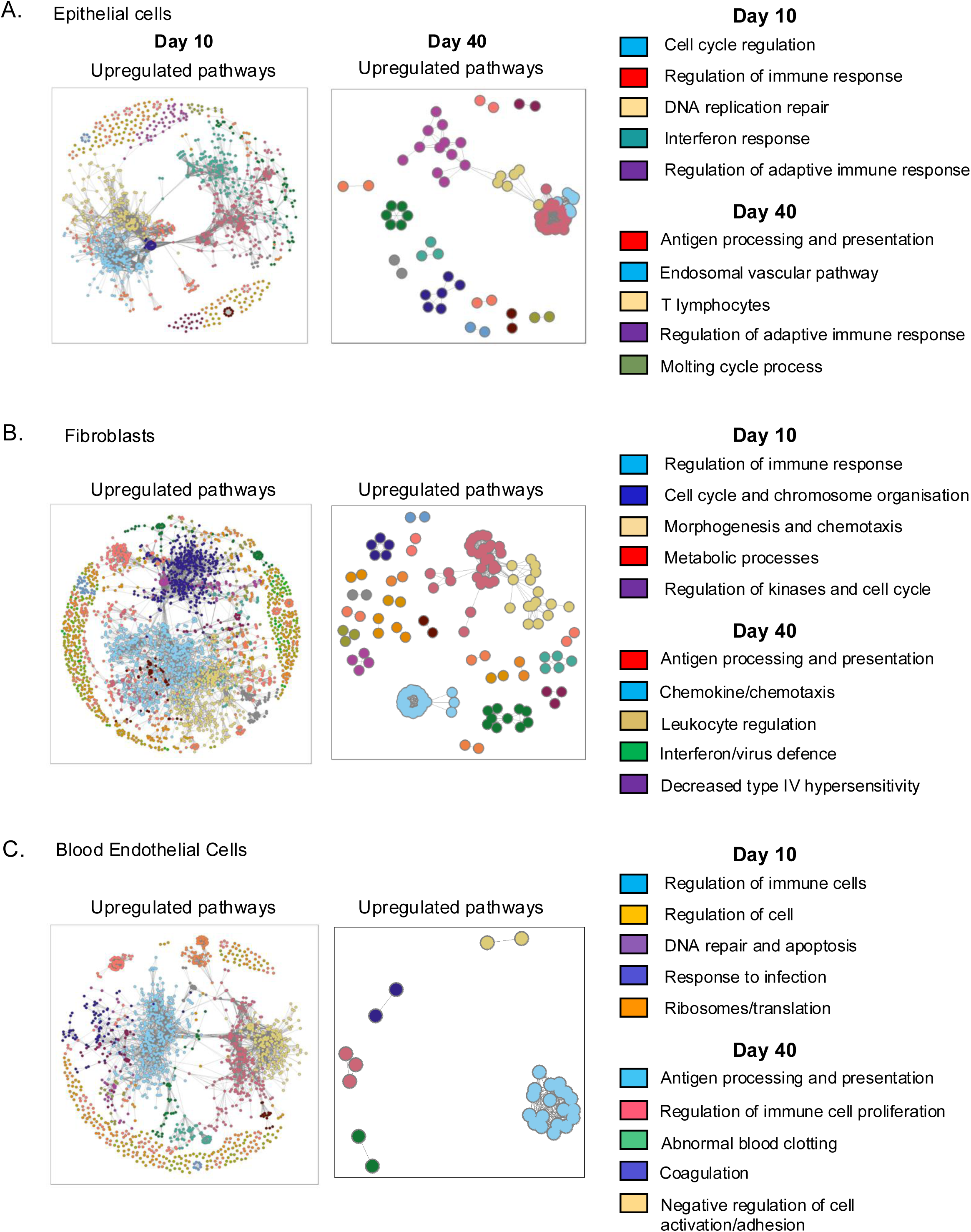
Network analysis shows conserved upregulation of genes involved in antigen processing and presentation in lung structural cells at day 40 post IAV infection. Networks of all significantly enriched genes sets in epithelial cells (A), fibroblasts (B), and blood endothelial cells (C), highlighting significance and gene set interconnectivity. Networks are given for enrichment amongst all significantly differentially expressed genes (p.adj < 0.05, absolute log2 fold > 0.5) between day 10 and naïve and day 40 and naïve. Enrichment analysis was performed using Hypergeometric Gene Set Enrichment on the gene set databases STRING11.5. For each network nodes represent gene sets and edges represent two gene sets with a Szymkiewicz-Simpson coefficient of at least 0.5.

### Expression of SpiB in airway epithelial cells found near immune cell clusters is dependent on viral replication

Analysis of the upstream regulators of the DEG identified at day 10 and 40 in epithelial cells and fibroblasts indicated a number of potential upstream regulators including ETS1, NFκB1 and CITED2 (SFig 4A). These molecules are downstream of Interferon (IFN)γ, CD40L and IFNα which can be produced by multiple cell types during IAV infection including epithelial cells, T cells and NK cells^31^. Interestingly, both epithelial cells and fibroblasts had significantly elevated gene expression levels of the ETS transcription family member *SpiB* at day 40 post infection and we confirmed increased expression by qPCR (SFig 4B). SpiB has recently been shown to co-ordinate differentiation and chemokine production by splenic fibroblasts following lymphocytic choriomeningitis virus (LCMV) infection^32^.

To investigate the potential contribution of SpiB to the transcriptional changes in lung structural cells following IAV infection, we compared the persistently upregulated DEG identified in our RNAseq analysis with genes identified as SpiB targets using a mouse ChIP-seq data set^33^. In epithelial cells, 25% of persistently upregulated DEG following IAV infection were SpiB target genes (20/80), in fibroblasts SpiB target genes (22/131) accounted for 17% of DEG (Fig 3A and Extended data 3). 27% of the SpiB targets induced by IAV infection were conserved between lung epithelial cells and fibroblasts (9/33): *H2-Q4*, *H2-Q5*, *H2-Q7, Psmb8*, *H2-K1*, *H2-Q6*, *SpiB*, *H2-Ab1*, *Psmb9* (Fig 3B). These data indicate that structural cells share an imprint of previous infection and SpiB may be a marker of this infection experience.

**Fig 3:**
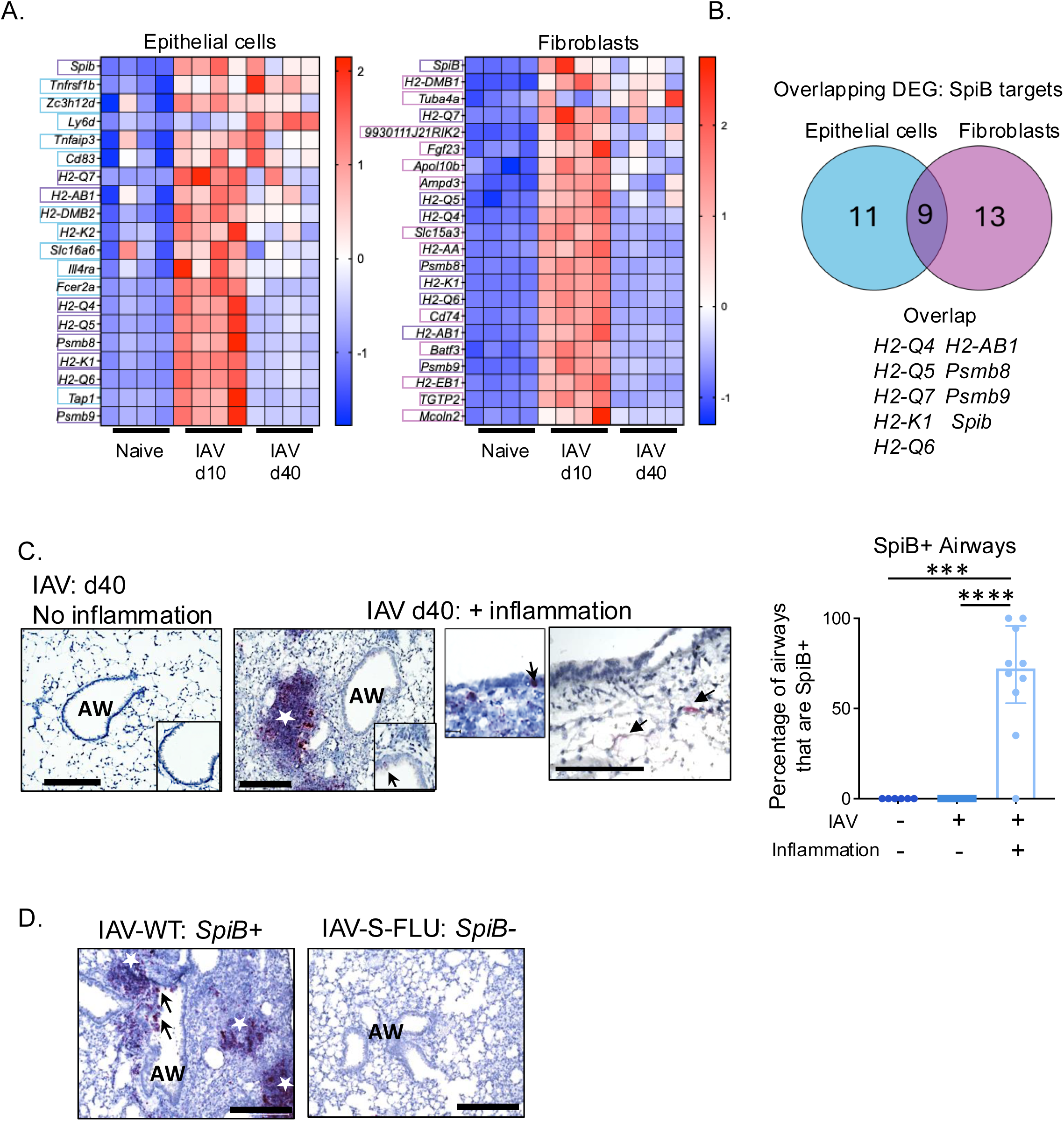
SpiB is upregulated in lung epithelial cells adjacent to areas of inflammation and this is dependent on viral replication. DEG upregulated at both day 10 and 40 post-infection from the experiments described in Fig 1 were compared with target genes of the transcription factor, SpiB, identified using a publicly available ChiPseq dataset (Solomon^33^ *et al*.) A: Heatmaps show SpiB target genes in epithelial cells and fibroblasts, respectively. Expression across each gene (or row) is scaled, with a Z-score computed for all DEG. Targets specific to epithelial cells (blue boxes), targets specific to fibroblasts (pink boxes) and those that overlap between cell types (purple boxes). Samples with relatively high expression of a given gene are marked in red and samples with relatively low expression are marked in blue. B: Venn diagram shows overlap between SpiB target genes and differentially expressed genes (DEG) in lung epithelial cells (blue) and fibroblasts (pink) post IAV infection. C: Lungs were taken from naïve or C57BL/6 mice infected with IAV 30-40 days previously. Infected lungs with no inflammation have *SpiB* negative airways while *SpiB* (black arrows) is localized in areas of inflammation (dark red color, indicated by white star) within the lung and in airways, close to areas of inflammation. Data are 4-6 mice per experimental group, combined from 2 independent experiments. Each symbol represents a mouse and bars show the median with the interquartile range as data are not normally distributed. Data analysed using Kruskal-Wallis with Dunn’s post hoc comparison test. ***: p <0.001 and ****: p <0.0001. D: C57BL/6 mice were infected intranasally with IAV (either wild type or single cycle (S-FLU), Airways (Aw), blood vessels (Bv) inflammatory foci/immune cell clusters (labelled with white stars) and *SpiB* positive airway epithelial cells (black arrows). All scale bars are 200µm.

To determine the location of transcriptionally altered epithelial cells and fibroblasts, we performed RNAscope on paraffin embedded lung tissue from naïve animals or mice infected 10 or 40 days previously with IAV. We found clusters of B220+ B cells that were SpiB+ cells in the parenchyma, as described by ourselves and others (SFig 5)^17,34–38^. SpiB+ epithelial cells were rare in the airway epithelium of naïve animals but were increased in SpiB+ structural cells in infected animals, but only present in areas of the lung close to clusters of immune cells (Fig 3C).

This link in location between infection altered epithelial cells and immune clusters could indicate sites of viral replication during the initial infection. To ask if viral replication was required for SpiB expression, we infected mice with a modified IAV virus, S-FLU, which, due to a mutation in the vRNA encoding HA, can only undergo one round of replication in host cells^39^. Despite this reduced replication, S-FLU does induce systemic and local adaptive immune responses^39,40^. *SpiB* expression was not detected in lung epithelial cells (or fibroblasts) in animals infected with S-FLU 30 days previously (Fig 3D). In addition, these mice did not display overt clinical signs of infection, with no reduction in total body weight following infection with S-FLU as previously reported^39^ (SFig 6A). At 30 days post infection lungs from mice infected with S-FLU had reduced levels of inflammatory infiltrate compared to those infected with the WT-IAV strain (SFig 6B) shown by H&E staining (SFig 6C). Viral replication was also necessary for the formation of immune cell clusters. These mice lacked B220+ and CD4+ cell containing clusters, compared to IAV-X31 infected mice (SFig 6D). Taken together, our findings indicate that viral replication is necessary for SpiB expression by lung epithelial cells.

### IAV infection leads to increased MHCII expression by multiple epithelial cell subsets

The lung contains multiple subsets of epithelial cells that are specialized for different roles and that respond distinctly to IAV infection^3,26,41^. We used multi-parameter flow cytometry to investigate the consequences of IAV on lung epithelial cell subsets. We coupled this with a novel SpiB reporter mouse to explore further the potential role for SpiB in epithelial cell function. In these transgenic (Tg) animals the sequence for a 2A peptide (T2A^42^) and the fluorescent molecule, mCherry, were inserted into the *SpiB* locus. As expected B cells^43^ and plasmacytoid DCs^44,45^ were positive for the reporter (SFig 7A). We also sorted SpiB-mCherry negative and positive EpCAM+ cells from naïve mice and examined the expression of *Spib* by semi-quantitative PCR (SFig 7B). SpiB-mCherry+ cells expressed low levels of *Spib* but higher levels than SpiB-mCherry negative cells. This low expression level likely explains why we do not detect *Spib*+ cells in naïve lungs by RNAscope. The SpiB-mCherry+ cells also expressed higher levels of SpiB target genes (*Psmb9, Tap1, Cd74, H2ab1, Tap1*^33^) and *Ifi47,* an Interferon-Stimulated Gene^46^ (SFig 7B).

In naïve mice, SpiB was expressed by multiple epithelial cell types including those of the upper airway: ciliated (CD24hi Sca1hi), club (CD24+ Sca1+), progenitor cells (CD24+ Sca1neg), Sca1+ progenitor cells (Sca1+ CD24neg), and alveolar epithelial cells (ATI: CD24neg Sca1neg Pdpn+ and ATII: CD24neg Sca1neg Pdpnneg MHCII+) from the distal lung (Fig 4A); gating strategy is shown in SFig 7C and based on previous studies^47,48^. In comparison to SpiB negative EpCAM1+ cells, the SpiB+ population was more likely to contain ATII cells but fewer ATI, and CD24+ and Sca1+ progenitor cells (Fig 4A).

**Figure 4:**
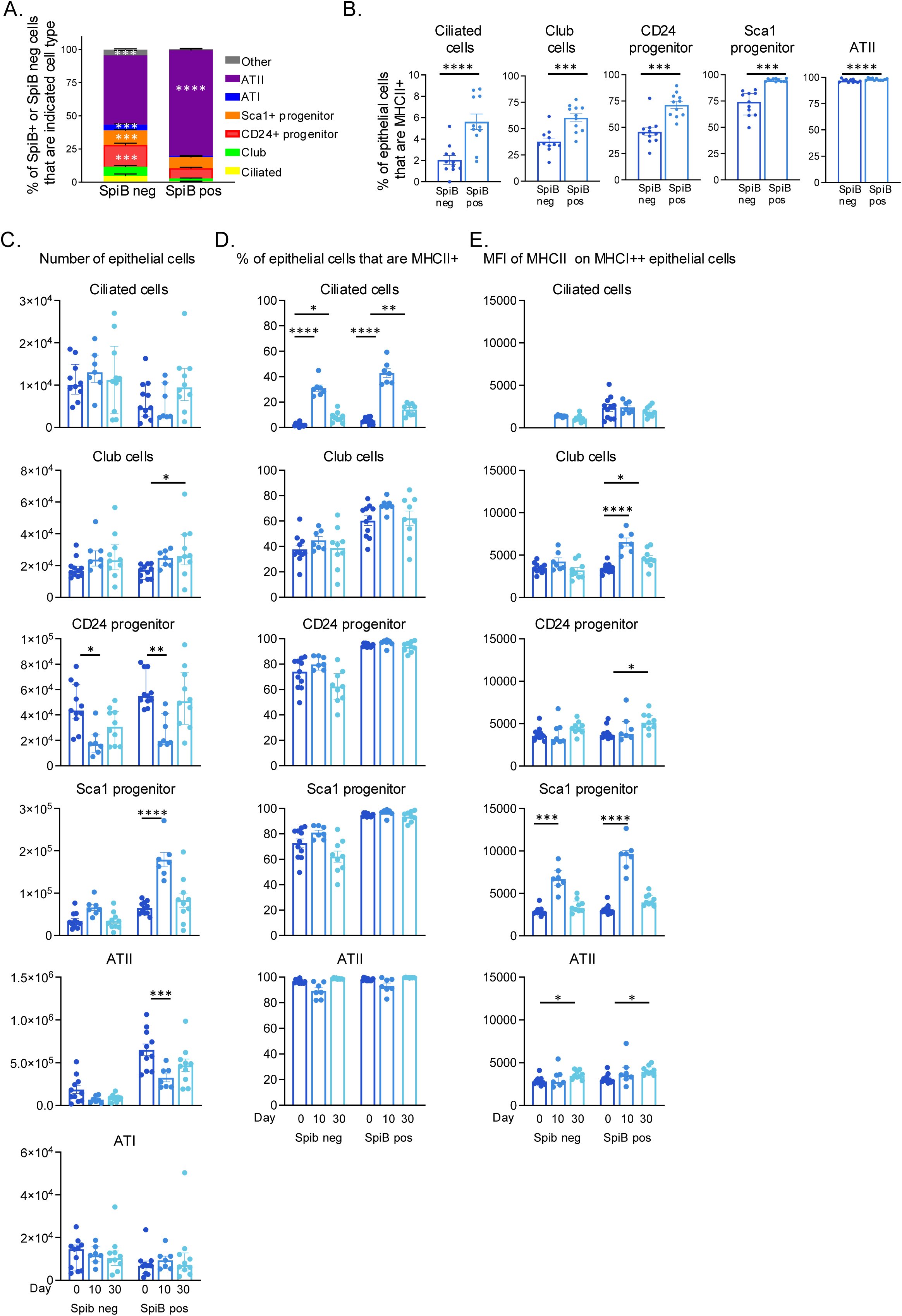
IAV infection results in upregulated MHCII expression by SpiB+ ciliated and progenitor cells at day 10. SpiB-mCherry reporter mice were infected with IAV (100-200PFU) and lungs taken at day 0, 10 or 30 post infection. EpCAM1+ epithelial cell populations were identified by gating on lineage negative (CD45, CD31 and CD140a neg), SpiB negative or SpiB+ populations. A: Frequency of SpiB positive or negative lung epithelial cells by cell subset in naive mice. B: Frequency of MHCII+ cells by each epithelial cell subset in naïve mice. C: Numbers of SpiB negative or positive populations at the different timepoints. D: Percentage and E: MFI of each population expressing MHCII at day 0, 10 and 30 post infection. In all experiments, data are combined with N=11 naïve, N=7 primary and N=10 memory from 2 independent experiments. In A data show mean with SEM and differences tested by ANOVA and Šidák’s multiple comparison test with significance stars located in the group in which the cell type is present at a higher frequency. In B-D: each symbol represents a mouse and the bars show means with SEM error bars for normally distributed data and median with interquartile range for non-normally distributed data. In B, all data are normally distributed apart from Sca1+ progenitor cells. In C, all data are not normally distributed apart from Sca1 progenitor and ATII cells. In D, all data are normally distributed apart from Sca1 progenitor cells. In E, all data are not normally distributed apart from ciliated and club cells. In B, normally distributed data are tested via paired T-tests and non-normal by a paired Wilcoxon test. In C-D, normally distributed data are tested via ANOVA with a Šidák’s multiple comparison test and non-normal by Kruskal-Wallis test followed by a Dunn’s multiple comparisons test. *:p <0.05, **:p<0.01,*** :p<0.001, ****: p<0.0001.

As SpiB is known to regulate molecules involved in MHCII antigen processing and presentation^33^ we examined the expression of MHCII on the epithelial cell populations, excluding ATI cells that are defined as MHCII negative^49^. For all the populations, SpiB+ cells were more likely to be MHCII+ than SpiB negative populations, this included ATII cells that are predominately MHCII+ in which we found that SpiB+ cells were slightly, but significantly, more likely to express MHCII (Fig 4B, SFig 7D).

We investigated the impact of IAV infection on the number of SpiB negative and positive epithelial cell populations at day 10 and 30 post-infection. IAV infection led to a decline of SpiB negative and positive CD24 progenitor cells and SpiB+ ATII cells at day 10 (Fig 4C). In contrast, Sca1+ SpiB+ progenitors more than doubled at day 10 post-infection while the numbers of SpiB negative Sca1+ progenitors remained unchanged. This increase in the SpiB+ population may reflect that Sca1+ progenitor cells can give rise to both bronchiolar and alveolar cells to repair the damaged epithelium^50^. All populations had returned to naïve levels by day 30 apart from a small increase in the number of SpiB+ club cells.

We also examined the expression of MHCII by the epithelial cells at the two infection time points. Ciliated cells were the most affected by the infection with both SpiB negative and positive populations more likely to express MHCII at day 10 and, while reduced, levels were still significantly increased at 30 post-infection compared to cells from naïve mice (Fig 4D). The amount of MHCII expressed by MHCII+ SpiB+ epithelial cells subsets was measured *via* MHCII geometric MFI (Fig 4E). SpiB negative and positive cells that were MHCII+ expressed broadly similar levels of MHCII. SpiB negative and positive Sca1+ progenitor cells expressed higher levels of MHCII at day 10 but not day 30 post-infection, compared to cells from naïve mice. In contrast, SpiB+ club cells expressed increased levels of MHCII at day 10 and these levels were still increased at day 30 post-infection. We also found small but significant increases of MHCII on CD24+ progenitor and ATII populations at day 30 post-infection compared to levels in cells from naïve mice.

Together, these data show that SpiB+ epithelial cells form a large proportion of the upper and distal airway epithelium and that while SpiB+ cells are more likely to express MHCII, SpiB is not required for MHCII expression. IAV infection had at most a minor lasting impact on the numbers of different epithelial cell subtypes but did lead to a sustained increase in the proportion of SpiB+ ciliated cells that expressed MHCII and increased levels of MHCII on several different epithelial cell subsets.

### Epithelial cells from IAV re-challenged mice have less virus than those from primary infected animals

We next investigated the consequences of a second IAV infection on lung epithelial cells and fibroblasts. To reduce the impact of neutralizing antibody responses, we infected mice with two different strains of IAV, WSN (H1N1) in the first infection and X31 (H3N2) in the re-challenge. We compared the response in the re-challenged mice with a primary X31 infection in age-matched naïve animals.

To identify infected epithelial cells and fibroblasts by flow cytometry we used a fluorescently labelled anti-IAV nucleoprotein (NP) antibody (Fig 5A). Mice infected 30 days earlier with WSN had reduced percentages and numbers of IAV-NP+ epithelial cells two days after infection with IAV-X31 compared to infected naïve mice (Fig 5A).

**Figure 5:**
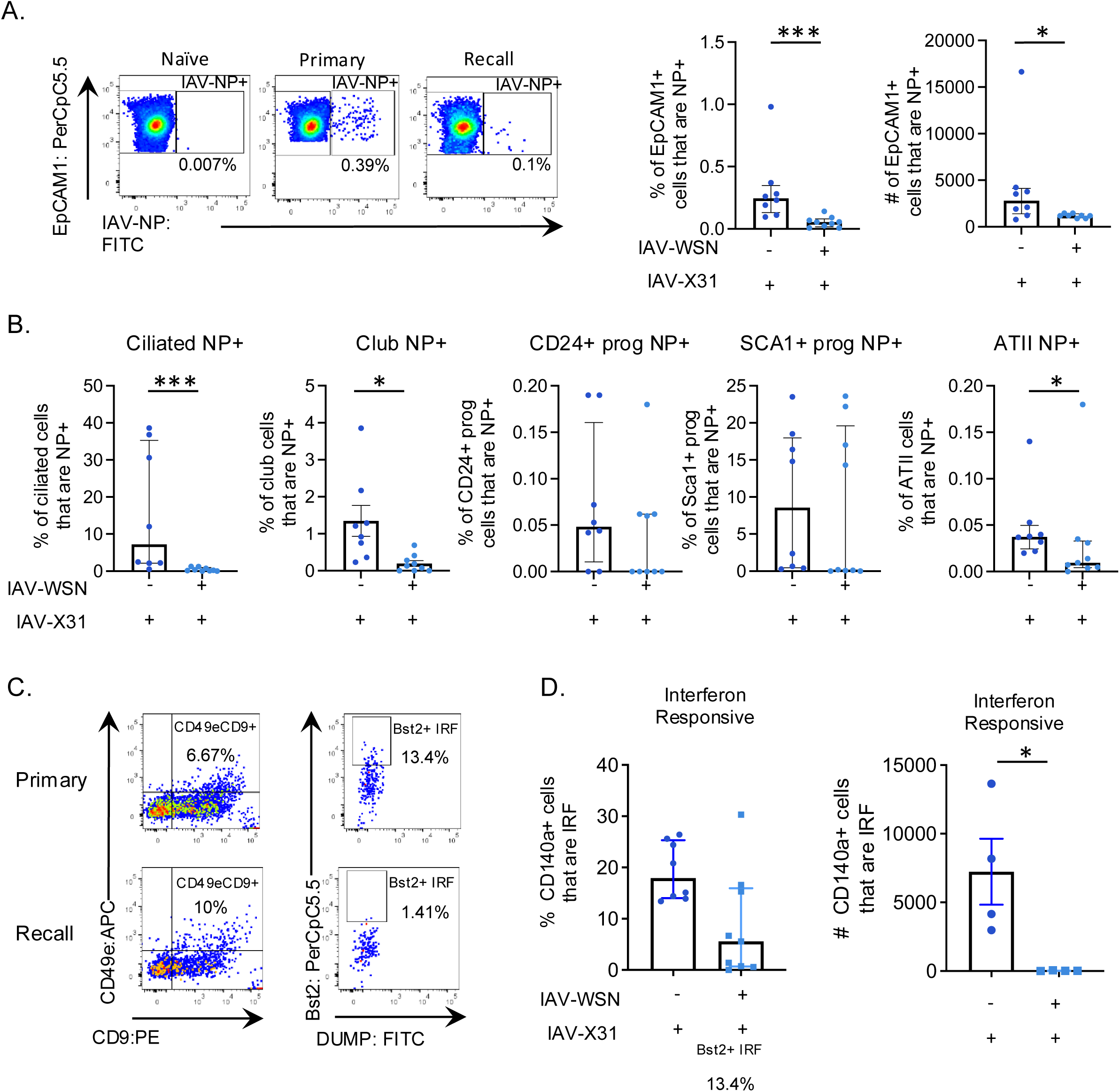
Epithelial cells from IAV re-challenged mice have less virus than those from primary infected animals. C57BL/6 mice that were either naïve or infected 30 days earlier with IAV-WSN, were infected with IAV-X31, their lungs harvested 2 days later, and epithelial cells and fibroblasts examined by flow cytometry. A: The percentages and number or EpCAM1+ cells positive for IAV-NP were examined by flow cytometry, gated as shown in SFig 7C. B: The percentages of each epithelial cell type that were IAV-NP+. C: Representative FACS plots of IFN-responsive fibroblasts indicating positive populations, cells are gated on live, single, dump negative (CD45/CD31/EpCAM1 negative) cells that are CD140a+ gated on CD49e+CD9+ fibroblasts that are Bst2+. D: the percentages and numbers of IFN-responsive fibroblasts. Data are from two independent experiments combined. In all graphs, each symbol represents a mouse, and the bars show means with SEM error bars for normally distributed data and median with interquartile range for non-normally distributed data. In A, data are not normally distributed. In B, all data are not normally distributed apart from Club cells. In D, the percentage of IFN responsive fibroblasts are not normally distributed and the number normally distributed. In all graphs, normally distributed data tested *via* t-test, and non-normally distributed data tested *via* Mann Whitney test. *: p <0.05, ***: p<0.001.

To examine this difference in more detail, we gated on the epithelial cell subsets and examined the percentages of these cells that were IAV-NP+ (Fig 5B). In primary infected animals, ciliated and Sca1+ progenitor cells were the most likely to be infected, followed by club cells. Very few CD24+ progenitor and ATII cells were infected and too few ATI cells were detected to examine. Prior infection led to reduced IAV-NP in ciliated, club, and a small decline in ATII cells, but no differences in the progenitor cells.

Few fibroblasts (CD140a+) were IAV-NP+ after IAV infection and there were too few cells detected to examine this further. The number of interferon-responsive^3^ fibroblasts were reduced in re-infected mice compared to primary infection (Fig 5C-D and gating in SFig 8). These data highlight that while fibroblasts are much less likely to be infected than epithelial cells, they are susceptible to local microenvironmental changes.

To determine if the reduction in IAV-NP levels by lung structural cells could be explained by enhanced/rapid phagocytosis of the virus by immune cells, IAV-NP levels were measured in lung myeloid and B cells. We found no evidence for this, all cells contained equivalent levels of NP apart from conventional dendritic cells 1 (cDC1) which contained less NP in the re-infected compared to the primary infected animals (SFig 9A-B).

### Re-challenge with IAV leads to rapid up regulation of genes involved in communication with T cells by lung epithelial cells

To investigate the consequences of re-infection in more depth, we compared gene expression in FACS sorted EpCAM1+ cells 2 days after a primary or re-challenge infection. EpCAM1+ cells from re-infected mice had increased expression of molecules in class I and II antigen presentation pathways (*B2m*, *H2-K1, H2-DMB2*, *Ciita*) and virus detection (*Nlrc5*) (Fig 6A-B and Extended data 4). Detection of IAV-NP+ cells requires fixation making it difficult to investigate gene expression in FACS sorted cells. Therefore, we investigated whether gene expression is different in infected cells in primary infected and re-infected animals by combining RNAscope with GeoMx spatial transcriptomics. First, we identified IAV-infected airways in primary and re-infected animals using a probe for IAV-X31 (Fig 6C). Only three of the five re-infected mice that we analyzed contained virus+ areas and we found a reduction in areas of lung containing virus in re-infected compared to primary infected animals (SFig 10A). We then examined gene expression by GeoMx in virus+ airway areas, and virus negative areas either in the same airway as the infected cells, or from adjacent airways (Extended data 5 and 6). We also included airways from sections of the lungs which did not contain any virus taken from primary and re-infected animals.

**Figure 6:**
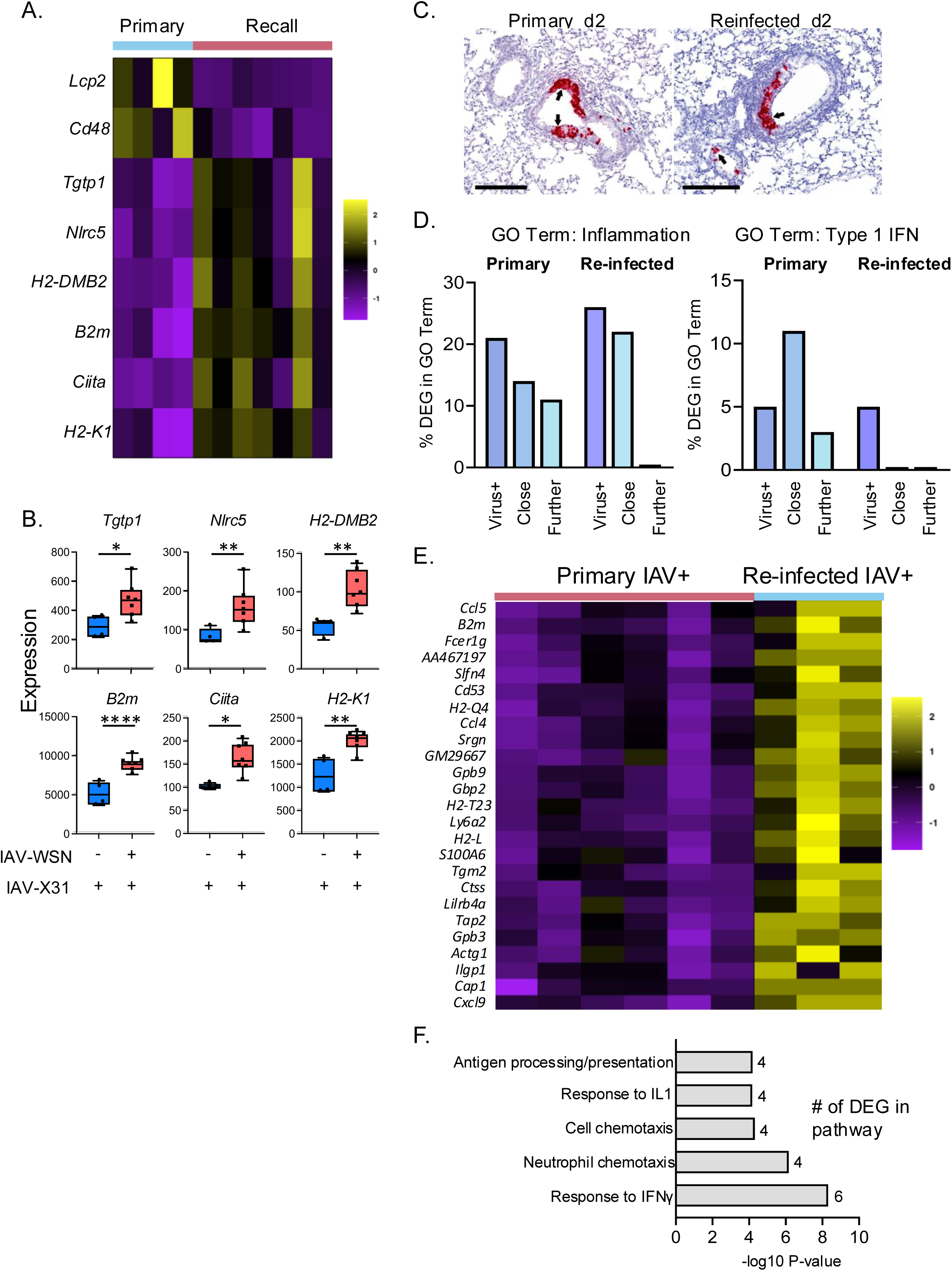
Spatial analysis of gene expression in primary and re-infected animals show altered inflammatory and type I IFN response and increased IFNγ response in re-infected animals. C57BL/6 mice that were either naïve or infected 30 days earlier with IAV-WSN, were infected with IAV-X31, their lungs harvested 2 days later, their epithelial cells were sorted by FACS and gene expression analyzed by Nanostring nCounter or lungs fixed and embedded in paraffin. A-B: Heatmap and example boxplots showing DEG from primary or re-infected animals in FACS sorted epithelial cells. DEG with a 2-fold change and an FDR q value < 0.05 in lung epithelial were considered statistically significant. Primary group (n=4) and re-challenge group (n=7), data are from two independent experiments combined and each dot represents a mouse. C: RNAscope analysis for X31-IAV was performed on FFPE lung tissue, representative X31-IAV+ areas are shown in red and labelled with black arrows to highlight virus + bronchial epithelium. Bars, 200 micrometres. Virus positive and negative airways from 6 primary and 3 re-infected mice were analysed by GeoMX. D: DEG between normal tissue and 3 areas: virus+ airway cells; virus negative cells within the same airway (Close); and adjacent virus negative airways (Further) were classified based on presence within MGI GO terms ‘Inflammation’ or ‘Type 1 IFN’. E: Heatmap showing DEG between X31-IAV positive airways in primary versus re-challenge groups, DEG with a 2-fold change and an FDR q-value < 0.05 in lung epithelial cells were considered statistically significant. F: Enriched pathways found in DEG identified between viral+ airways in day 2 primary or re-infected C57BL/6 mice analysed by GeoMx. The x-axis represents the −log10 of the P value given during the analysis; (FDR < 0.05) with the number of DEGs listed from each pathway.

As expected, in primary infected mice a comparison of gene expression in virus+ areas versus airways from non-infected sections demonstrated an upregulation of anti-viral genes including *Cxcl10*, *Isg15* and *Irf7* and upregulation of pathways related to ‘defense to viruses’ and ‘type 1 IFN’ (SFig 10B, Extended data 7). An anti-viral response was also evident in virus negative areas within infected airways and adjacent virus negative airways. Similar to the primary infected mice, in re-infected animals, genes within ‘type 1 IFN’ and ‘response to virus pathways’ were upregulated in virus+ airways compared to airway cells from non-infected sections from the same mice (SFig 10C, Extended data 7). To compare the responses between primary and re-infected animals, we performed two types of analysis. First, we counted the number of upregulated genes within the Gene Ontogeny (GO) terms ‘Inflammation’ and ‘Type 1 IFN’ identified in virus+, virus negative areas within virus+ airways (close), and virus negative adjacent airways (further) and calculated these as a percentage of the total DEG within each area (Fig 6D, Extended data 7).

All three areas had genes within both GO terms in primary infected animals, although ‘further’ areas had the lowest number. In contrast, in re-infected animals while genes within GO term ‘Inflammation’ were upregulated in virus+ and ‘close’ airways, no genes in this term were found in ‘further’ areas and no genes in GO term ‘Type 1 IFN’ were found in ‘close’ or ‘further’ areas in re-infected animals. These data suggest that while IAV may induce inflammatory type 1 IFN responses in infected cells in primary and re-infected mice, the response is much more contained in re-infected animals.

Second, we compared gene expression between virus+ airways taken from primary or re-infected animals (Fig 6E, Extended data 7). We found increased expression of genes in the GO term ‘Cellular Response to IFNγ’ including *Ccl5* and *Cxcl9* in the airways of re-infected mice (Fig 6F). These data suggest that IFN-γ producing immune cells, including CD4 and CD8 T cells^51^ act on infected cells that in turn upregulate molecules that enhance communication between immune and infected structural cells.

### Infection experienced structural cells more effectively control IAV than cells from naïve mice

The potentially increased communication between infected airway cells and T cells indicated that at this early timepoint following infection, the enhanced protection provided by a prior IAV-infection will be dependent on T cells. To test this, we infected two cohorts of C57BL/6 mice with IAV-WSN and treated one cohort with IgG and the second with depleting CD4 and CD8 antibodies (SFig 11A). Mice were treated 2 days prior and on the day of the infection with X31; control experiments demonstrated a loss of CD4 and CD8 T cells in lymphoid organs and the lung at day 2 post-re-infection (SFig 11B). In data consistent with Epstein *et al*^29^., we found that mice treated with IgG and T cell depleting antibodies had less virus in their lungs compared to primary infected animals (Fig 7A). Moreover, there was no difference in levels of virus between the two re-infected groups. These data suggest that T cells are not required for enhanced protection at day 2 post-infection and that, potentially, enhanced viral control responses are from the infected cells themselves.

**Figure 7:**
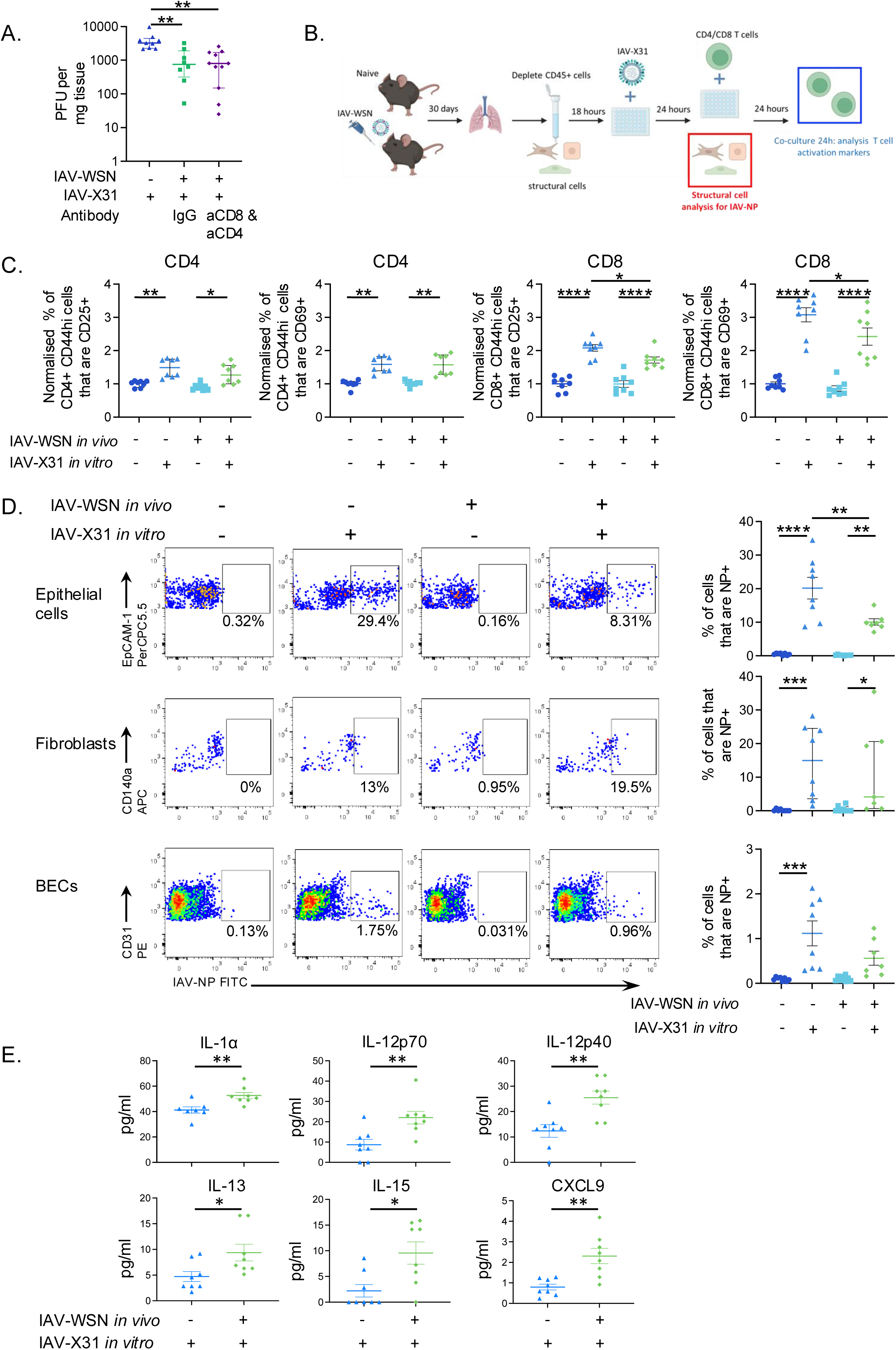
Infection experienced stromal cells display enhanced viral control and can reactivate T cells. A: C57BL/6 mice were infected with IAV-WSN on day 0 and treated with 400μg isotype control or 200μg each of anti-CD4 (GK1.5) and anti-CD8 (2.43) on days 28 and day 30 when mice were (re)-infected with IAV-X31, combined from two experiments: Primary (n=9) mice, recall IgG (n=9), recall aCD4aCD8 (n=11). B: Schematic showing experimental design of IAV infection and *in vitro* re-challenge and co-culture. Lung CD45 negative cells were isolated from naïve or C57BL/6 mice infected 30 days earlier with IAV-WSN. After 24hours, the cells were infected with IAV-X31 and the cells examined by flow cytometry after a further 24hours or co-cultured with T cells isolated from the spleens of mice infected with IAV-X31 9 days earlier. C: The percentage of CD44hi CD4 or CD8+ T cells that were CD25 or CD69 positive, normalized to the mean of the no infection control within each of the 2 experiments with 4 mice within each naïve and IAV-memory group per experiment. D: Representative FACS plots and data for IAV-NP for each cell type, graphed data are average of two technical replicates. Cells were gated on live, single, CD45 negative EpCAM1+ (epithelial cells), CD140+ (fibroblasts), and CD31+ (blood endothelial cells). Numbers in plots indicate the percentages of cells that are IAV-NP+. E: Supernatants from the infected cultures were tested for multiple immune mediators by Luminex 24 hours after *in vitro* infection. Data in D-E combined from the same two experiments, Primary: n=8, Re-infection: and n=7. In all graphs, each symbol represents a mouse and the bars show means+/-SEM for normally distributed data and median with interquartile range for non-normally distributed data. In A, data are not normally distributed. In C, CD4 data are not normally distributed and CD8 data normally distributed. In D, epithelial cell and BEC data are normally distributed, and fibroblast data are not normally distributed. In E, all data are normally distributed. In A-D, non-normally distributed data tested with a Kruskal-Wallis test followed by a Dunn’s multiple comparisons test. In C-D, normally distributed data tested by ANOVA followed by a Šidák’s multiple comparison test. In E, data tested by t-test. *:p<0.05,**: p<0.001,***: p <0.001, ****: p<0.0001.

We chose to use a reductionist *in vitro* model to investigate this finding further, reducing the complexity of a rapidly changing response *in vivo* (Fig 7B). First, we asked whether the prior infection altered the ability of the structural cells to present antigen to CD4 and CD8 T cells. To achieve this, we isolated CD45 negative cells from naïve and day 30 IAV-WSN infected animals and rested these overnight before infecting them with IAV-X31 *in vitro*. The following day, T cells were isolated from the spleens of mice infected with IAV (X31; day 9/10 post infection) to generate a pool of IAV specific CD4 and CD8 T cells. These cells were co-cultured with the infected CD45 negative cells and expression of T cell activation markers, CD25 and CD69, was analyzed by flow cytometry after a further 24 hours. CD4 and CD8 T cells co-cultured with *in vitro* IAV-infected CD45negative cells from either naïve mice or previously infected mice expressed higher levels of CD25 and CD69 than T cells co-cultured with control CD45 negative cells that did not receive IAV *in vitro*. This demonstrates that the structural cells could process and present IAV antigens to CD4 and CD8 T cells (Fig 7C and SFig 12A-B).

We expected that CD45 negative cells from previously infected animals might present antigens more effectively to T cells given their increased expression of molecules involved in these processes. Expression of CD25 and CD69 were equivalent between CD4 T cells cultured with IAV-infected structural cells taken from naïve or IAV-infected mice. There was a slight but significantly lower expression of CD25 and CD69 on CD8 T cells cultured with IAV-X31 infected structural cells from previously infected animals compared to X31-infected structural cells from naïve mice.

Potentially, prior infection enabled the CD45 negative cells to control the virus more effectively and thus these cells might contain lower levels of antigen. To test this, we harvested a separate plate of structural cells 24 hours after *in vitro* infection and examined the levels of IAV-NP in epithelial cells, fibroblasts and BECs by flow cytometry. Few BECs were infected with IAV, in contrast, clear populations of fibroblasts and epithelial cells were IAV-NP+ (Fig 7D). More NP+ cells were found in infected cells compared to the no infected controls. The exception was BECs from previously infected mice in which there was no significant difference between the no infection and infection control, suggesting these cells may be controlling the virus more effectively than cells from naïve animals. However, there was also no significant difference between the two infected groups. In contrast, there were twice as many NP+ epithelial cells in cultures from naïve mice compared to cells from previously infected animals. In the fibroblasts, there was no difference in the percentages of NP+ cells between the infected groups. These data suggest a prior IAV infection enhances the ability of epithelial cells to control a subsequent challenge and demonstrate a cell intrinsic effect as fibroblasts within the same cultures were infected at the same rate, regardless of the cell source.

To ask whether this reduced infection led to a lower inflammatory response, we examined culture supernatants for the presence of cytokines and chemokines 24h after the *in vitro* infection using a multiplex array. The supernatants from CD45 negative cells isolated from previously infected mice, contained higher levels of IL-1α, IL-12 p40 and p70, IL-13, IL-15, and CXCL9 compared to supernatants from cultures with cells from naïve mice (Fig 7E). Type I and III IFN were not detectable by ELISA (below the limit of detection; 31.3 pg/ml). In conclusion, these data suggest that the rapid control of virus by the structural cells is accompanied by an increased inflammatory cytokine response, perhaps to warn and attract immune cells to control any virus not contained by the epithelial cells themselves.

## Discussion

Our functional and geographical analysis of the post-IAV lung demonstrates substantial and sustained changes to the three main structural lung cell types, epithelial cells, fibroblasts, and blood endothelial cells. All three cell types maintain upregulated expression of genes involved in antigen processing and presentation, suggesting that they may have an enhanced ability to communicate with T cells. While interactions between T cells and lung structural cells may be important^52,53^, we found that lung epithelial cells from mice previously infected with IAV can themselves control IAV more efficiently than cells from naïve mice. These data demonstrate enhanced protective responses by epithelial cells indicating these cells display protective immune memory.

Evidence for innate or trained immunity in multiple cell types has been growing over the last ten years^8,24,54^. Skin epithelial cells previously exposed to inflammatory stimuli respond more rapidly through an inflammasome-dependent mechanism^55^ and lung epithelial cells that survive an IAV infection have altered gene expression and inflammatory responses following re-infection^25–27^. Fibroblasts are known to play important roles in lung anti-viral responses and in other organs, including synovial fibroblasts in joints and gingival fibroblasts, can display evidence of memory to previous insults^54,56–59^. Most of the evidence for sustained changes in BECs come from the field of cardiovascular disease where chronic inflammation may sustain altered responses^54,60–62^. Our data extended the field by examining all three cell types within one organ, demonstrating a shared sustained response in all cell types suggestive of coordinated tissue memory.

Epithelial cells, most notably ATII cells can express MHCII^52,53,63^. Antigen presentation by lung epithelial cells is required for normal responses to *Streptococcus pneumoniae* and Sendai virus infections in mice^52,53^. We found that IAV infection led to upregulation of MHCII either by the proportion of cells that were MHCII+ or increased expression on upper (ciliated, club), progenitor (CD24+, Sca1+), and lower airway (ATII) epithelial cells at day 10 post-infection. Some of these changes, including more ciliated cells expressing MHCII, persisted at day 30 post-infection. As ciliated cells can live for months^64^, our findings may indicate these MHCII+ ciliated cells persist from the infection, while other changes may be sustained by the progenitor epithelial cells.

Indeed, Fiege *et al*., demonstrated that ciliated epithelial cells can survive IAV infection^27^. Interestingly, other mouse models of epithelial lung injury^65^ also reveal upregulation of MHCII by ciliated cells and a recent human challenge study, found a small subset of ciliated cells hyper-infected with SARS-CoV2 that were MHCII+ but that expressed anti-inflammatory molecules^66^. This study identified HLA-DQA2 (associated with MHCII antigen presentation) as a predictor of infection outcome and protection^66^. Sustained elevated MHCII levels may reflect an imprint of infection, providing these epithelial cells with an advantage by more quickly altering T cells to a new infection. However, as non-professional APCs, the cells could drive T cell tolerance rather than activation^67^

We bioinformatically identified the transcription factor, SpiB, as a potential upstream regulator of the altered MHCII expression and upregulation of gene expression in multiple molecules involved in antigen processing and presentation^33^. In addition to B cells, SpiB is expressed by plasmacytoid dendritic cells^45,44^, Micro-fold cells^28^, tuft cells, some fibroblasts in secondary lymphoid organs^68^, and some thymic epithelial cells^69^. Our data suggest that it is also expressed by multiple populations of lung epithelial cells and that SpiB+ positive cells express higher levels of MHCII. Recently, two studies identified SpiB+ epithelial cells in distinct anatomical locations. Surve *et al*., performed re-analysis of publicly available single-cell RNA sequencing data and detected rare M cells in the homeostatic mouse trachea^70^. While Barr *et al*., used single-nucleus RNAseq to detect lung M cells expressing *Tnfaip2*, *Sox8*, *Spib*, *Ccl9*, *Ccl20*, and *Tnfrsf11a* ^71^.

We showed that following IAV infection, *Spib*+ upper airway epithelial cells were likely to be close to persistent clusters of immune cells. These observations are consistent with the study by Barr *et al*., where *Spib*+ cells in the IAV infected lung persisted from day 5 to 56 post infection^71^. However, our study provides additional insight by validating this at the protein level and by examining MHCII expressing in naïve and infected animals. We and others have characterized these clusters which predominately contain B and T cells^17,34–38^. The close proximity of these immune cells suggests that they may release molecules to sustain the altered epithelial cells, and/or their proximity evidences the past location of viral replication.

Our data support that fibroblasts can contribute to the early anti-viral response through multiple mechanisms, including communication with T cells. Although fibroblasts are much less likely to become infected with IAV than epithelial cells^10^, several studies have highlighted their ability to drive immunopathology or restrain lung inflammation by communicating with lung T cells^72,73,74^. Boyd *et al*., showed that damage responsive fibroblasts can influence the migration of CD8 T cells that enter inflamed tissues and that this amplifies immunopathology^3^. Conversely, amphiregulin producing regulatory T cells promote alveolar epithelial regeneration and repair by activating Col14+ adventitial fibroblasts, supporting progenitor cells after IAV-induced damage^72^. Conditional deletion of these Col14+ fibroblasts reduced survival rates following IAV infection. Additionally, Treg cells secrete IL-1Ra, which suppresses IL-1 receptor-mediated chemokine production by adventitial fibroblasts, mitigating excessive inflammation^73^; disruption of this regulatory axis caused exacerbated inflammation and impaired viral control. Our investigation adds further complexity, suggesting that previous viral infections could alter the consequences of T cell-fibroblast interactions characterise by these studies of primary infections.

A classic characteristic of immune memory is the ability to respond rapidly to re-infection^75^. By measuring gene expression and developing an *in vitro* infection model, we demonstrated that lung epithelial cells display an enhanced response to re-infection leading to more rapid control of IAV.

This enhanced functional response was accompanied by increased production of cytokines. In primary infected mice, Hamele *et al.,*^76^ identified a population of IAV infected ciliated cells that produced the majority of inflammatory cytokines early in the response. Surviving ciliated cells may not only therefore, express more MHCII, but may also be responsible for the enhanced cytokine response we found in *in vitro* infection of CD45 negative cells from previously infected mice.

Krausgruber *et al.,* found that even in naïve animals, structural cells are poised, ready to produce molecules more often associated with immune cells^77^. Our data extend this finding, suggesting that exposure to pathogens boosts this homeostatic immune gene expression making structural cells an even more important component to protective immunity. Our study also provides rationale for attempting to target structural protective immunity as part of vaccine design, for example, by adjuvants. This concept was investigated in a recent study by Denton *et al*., who found that targeting TLR4 using adjuvants improved the MAdCAM-1 + stromal cell response to immunization and facilitated longer term protection post vaccination^78^.

Single cell RNAseq experiments examining primary IAV-infection have identified that ciliated and club cells can exhibit transcriptional signatures of interferon signalling independent of infection but dependent on the inflammatory microenvironment^76,79,80^. Similarly, our spatial transcriptomic analysis (GeoMx) of mouse lungs from primary infected mice, identified that genes associated with the type I IFN response and inflammation were found beyond virus+ regions. These responses were more restricted in lungs from re-infected mice in which the anti-viral response was contained close to the virus. This potentially limits inflammation driven immunopathology and the release of cytokines that cause systemic symptoms associated with the infection.

Our data show that early protection to re-infection was independent of T cells, but using plaque assays and RNAscope, virus is still present at day 2 following the re-challenge infection. Both CD4 and CD8 T cells protect mice and humans from IAV re-infection, especially when priming and re-challenge strains express different hemagglutinin and neuraminidase proteins^18,20,21,29,30,81,82^. The sustained expression of inflammatory chemokines and molecules involved in the processing and presentation of antigen following infection suggests that lung structural cells will have an enhanced ability to attract and communicate with local T cells. In summary, therefore, our data suggests that lung epithelial cells may play both early T cell independent and then later T cell dependent roles in accelerating viral clearance following re-infection.

## Materials and Methods

### Animals

SpiB-mCherry reporter mice were generated at MRC Harwell. The sequence for a 2A peptide (T2A^42^) with a GSG linker and the fluorescent molecule, mCherry, was inserted into the SpiB locus: Ensembl ENSMUSG00000008193. The knock in was inserted at the end of the coding sequence. SpiB reporter mice were viable, fertile and born at expected Mendelian ratios.

Ten-week-old female C57BL/6 mice were purchased from Envigo (United Kingdom), and male and female SpiB reporter mice were bred and housed at the University of Glasgow under specific pathogen-free conditions in accordance with UK home office regulations (Project Licenses P2F28B003 and PP1902420) and approved by the local ethics committee. No formal randomization was performed, and sample size was based on previous studies.

### IAV infections

SpiB or C57BL/6 mice were briefly anesthetized using inhaled isoflurane and infected with 100–200 plaque-forming units (PFU) of IAV WSN strain, depending on their age, weight, and sex or 200PFU IAV-X31 strain in 20 µL of phosphate-buffered saline (PBS) i.n. IAV was prepared and titered in Madin-Darby Canine Kidney cells (MDCK). Infected mice were weighed and any animals that lost more than 20% of their starting weight were humanely euthanized. Cages with mice that lost weight were given soft diet until all mice returned to their starting weight or above.

### S FLU virus

S-FLU virus^39^ was obtained from the laboratory of Alain Townsend at the university of Oxford. C57BL/6 were infected with 5 HAU (hemagglutinating units) of either the parental strain of the virus IAV-WT X31 or the S-FLU-X31.

### T cell depletion experiment

Two cohorts of C57BL/6 mice were infected with IAV-WSN and one cohort injected with 400µg of rat IgG (LTF-2) and the second with depleting 200µg CD4 (GK1.5) and 200µg CD8 (2.43) antibodies, all antibodies from BioXCell. Mice were treated intraperitoneally with anti-CD4/CD8 2 days prior to re-challenge with IAV-X31 and 2 days following infection. Mice were culled 2 days post re-challenge. Control IgG and anti-CD4/CD8 treatments were given to mice within the same cages to prevent cage-specific effects acting as confounders.

### Plaque assay on mouse lung homogenates

Once euthanized, mouse lungs were extracted and frozen at −80 °C. Lungs were thawed, weighed and homogenized with 1 ml of DMEM. Homogenates were cleared by centrifugation and titrated by plaque assay. Confluent monolayers of MDCK cells were washed once with PBS, infected with serial dilutions of virus and incubated for 1 h at 37 °C to allow virus adsorption to the cells. After virus removal, cells were overlaid with MEM with oxoid agar and incubated for 2 days at 37°C, 5% CO2. After removing the overlay, cells were fixed with PBS:4% formaldehyde and stained with a 0.08% Crystal Violet solution in 6.25% methanol solution for at least 1 h. Staining solution was rinsed under tap water, plates were air-dried, plaques were then counted by a blinded researcher.

### Tissue preparation

Mice were euthanized by cervical dislocation (RNA-sequencing, FACS sorting and flow cytometry analysis) or alternatively with a rising concentration of carbon dioxide (lung histology and imaging). For flow cytometry analysis of structural cells, single cell suspensions of lungs were prepared by enzymatic digestion with a final concentration of 1.6 mg/mL Dispase, 0.2 mg/mL collagenase P (Roche, UK) and 0.1 mg/mL DNase (Sigma, UK) for 40 minutes at 37°C in a shaking incubator and tissues disrupted by passing through a 100μm filter. Red blood cells were lysed with lysis buffer (ThermoFisher).

For analysis of CD4 and CD8 T cells by flow cytometry prior to euthanasia by cervical dislocation, mice were injected intravenously with 1μg AF488-conjugated anti-CD45 (clone, 30F11, ThermoFisher) and organs harvested after 3 minutes. Single-cell suspensions of lungs were prepared by digestion of snipped lung tissue with 1 mg/mL collagenase and 30μg/mL DNase (Sigma) for 40 minutes at 37 °C in a shaking incubator and tissues disrupted by passing through a 100μm filter. Spleens and lymph nodes were processed by mechanical disruption. Red blood cells were lysed from spleen and lungs with lysis buffer (ThermoFisher). Cells were counted using a hemocytometer with dead cells excluded using Trypan Blue.

### Flow Cytometry Staining

Cells were incubated for 10 mins with Fc block (homemade containing 24G2 supernatant and mouse serum) surface stained with anti-CD45 FITC (Biolegend, 30-F11) or anti-CD45-PE (BD, 30-F.11) or anti-CD45 eflour450 (ThermoFisher, 30-F.11), anti-CD31 PE (Biolegend, MEC13.3) or anti-CD31-FITC (BioLegend, MEC13.3), CD326/EpCAM1-1 PerCP-Cy5.5 (Biolegend, G8.8) or CD326/EpCAM1-1 BV711 (Biolegend, G8.8), anti-CD140a/Pdgfra APC (Biolegend, APA5) or anti-CD140a/Pdgfra BV605 (Biolegend, APA5), Gp38/podoplanin PeCy7 (Biolegend, 8.8.1), anti-Sca1 APC-Cy7 (BioLegend, W18174A), anti-CD24 BV421 (Biolegend, M1/69), anti-MHCII BUV395 (BD: 2G9), anti-Siglec F-APC (BioLegend, S17007L), anti-CD11b-PeCy7 (BioLegend, M1/70), anti-Ly6G-BV785 (BioLegend, 1A8), anti-Ly6C-PerCP-Cy5.5 (ThermoFisher, HK1.4), anti-CD64-BV711 (BioLegend, X54-5/7.1), anti-CD11c-eFluor780 (ThermoFisher, N418), anti-B220-eFluor 450 (RA3-6B2, ThermoFisher), anti-CD103-Qdot 605 (BioLegend, 2E7), anti-Bst2-PerCp-C5 (BioLegend, 927) and anti-CD9-PE (BioLegend, MZ3). Cells were stained with a fixable viability dye eFluor 780 or eFluor 506 (both ThermoFisher) as per the manufacturer’s recommendations. Cells were fixed with cytofix/cytoperm (BD Bioscience) when required for 20 min at 4°C and stained in permwash buffer with anti-IAV-NP FITC (Invitrogen, D67J) for 1 h at room temperature.

For T cells, surface stains were anti-CD4 APC-Alexa647 (RM4-5, ThermoFisher), anti-CD8 BUV805 (BD 53-6.7), anti-CD44 BUV395 (BD, IM7), anti-CD25 BV711 (Biolegend, PC61), anti-CD69 PE (BD, H1.2F3) anti-B220 eFluor 450 (RA3-6B2), anti-MHCII eFluor 450 (M5114) and anti-F480 eFluor 450 (BM8) all ThermoFisher. Anti-B220, MHCII and F480 were used as a ‘dump’ gate. Samples were acquired on a BD Fortessa and analyzed using FlowJo (version 10 BD Bioscience, USA).

### Lung digestion and isolation of mouse lung structural cells by Fluorescence-Activated Cell sorting

For FACS isolation of lung structural cells, single cell suspensions were generated by digesting snipped lungs with a final concentration of 3.2mg/mL Dispase-II, 0.4mg/mL Collagenase P, and 0.2mg/mL Dnase-1. Lung samples were incubated at 37°C for 20 minutes. Red blood cells were lysed using RBC buffer (ThermoFisher). Cells were simultaneously stained with CD45 microbeads (Miltenyi Biotec) and antibodies: anti-CD45 FITC (Biolegend, clone:30-F11) anti-Ter119-FITC (BioLegend, TER-119), anti-CD31 PE (BioLegend, MEC13.3), CD326/Epcam-1 PerCP Cy5.5 (Biolegend, G8.8), CD140a/Pdgfra APC (Biolegend, APA5). Hematopoietic cells were depleted using the LS MACS (Miltenyi Biotec) column as per manufacturer’s instructions. CD45 depleted lung cells were stained with eFlour 780 viability stain (Thermofisher), and cells sorted on BD FACS Aria IIU sorter. Cells were either sorted directly into RLT or first sorted into 50% FCS/RPMI, washed with PBS and then lysed in RLT buffer.

### RNA extraction for qPCR and nanostring analysis

RLT suspensions were centrifuged through Qiagen QIA-Shredders and RNA was extracted from sorted cells using the RNeasy mini kit (Qiagen,UK) with the following changes to manufacturer’s instruction: input to the RNeasy column was at a ratio of 100µL sorted cells: 350µL RLT: 200µL 100% EtOH. Following extraction samples were DNase-1 treated. For qPCR analysis, briefly, the first strand cDNA synthesis was conducted using a Reverse Transcription System kit according to the instructions of the manufacturer (Promega, Madison, WI).

### qPCR analysis

Gene expression was analyzed by qPCR (SYBR Green FastMix (Quanta Bioscience) on a QuantStudio 7 flex and expression was calculated using standard curves and results normalized to *Gapdh* or *18s* expression. qPCR standards were prepared using spleen or lung cells from IAV infected mice and purified by gel extraction (Quick Gel Extraction kit, Invitrogen) or PureLink PCR Purification kit (Invitrogen).

**Table 1:**
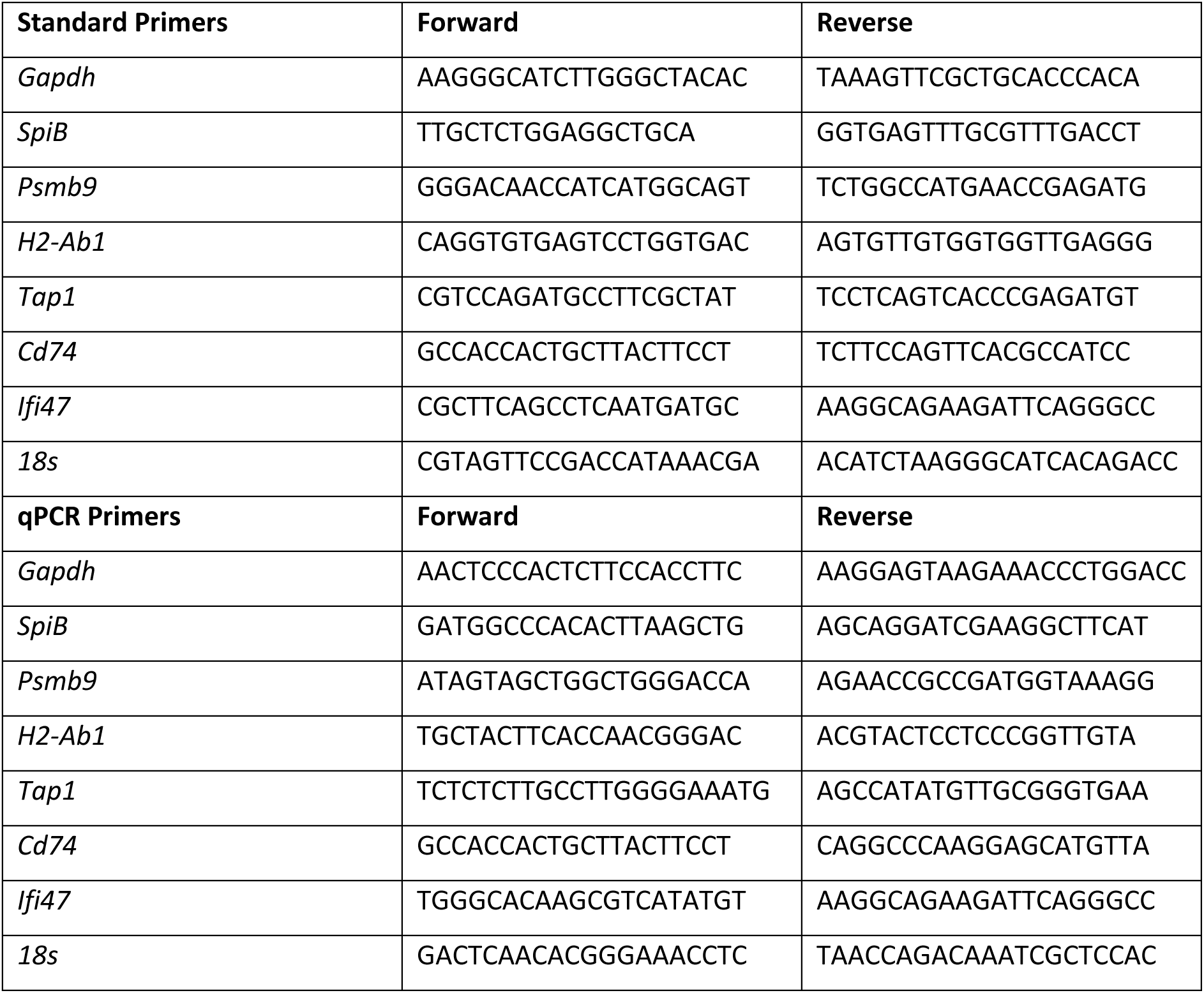
Primer sequences.

### NanoString nCounter analysis of sorted epithelial cells

Briefly, RNA from sorted lung epithelial cells of either naïve or IAV infected mice (day 30) was analyzed using the nCounter Mouse Immunology Panel (catalog # 115000052 XT_PGX_MmV1_immunology). RNA was purified using an RNA Concentration and Clean-Up kit as per manufacturer’s instruction (Zymo, Cambridge Biosciences, R1013). The integrity and quantity of total RNA were determined using Agilent 2100 Bioanalyzer (Agilent Technologies), samples with a RIN value of ≥8 were included for downstream analysis. This panel includes 547 genes covering core pathways and processes of the immune response, and 14 internal reference genes, a custom code set of 29 genes was used in conjunction with this panel (Extended data 2). Each sample consisted of RNA from sorted epithelial cells from one individual mouse. Assays were performed using 50ng input RNA and quantified on the nCounter FLEX system, sample preparation station, and digital analyzer (NanoString Technologies) according to the manufacturer’s instructions. Briefly, reporter and capture probes were hybridized to target analytes for 20 h at 65°C as per^68^. After hybridization, samples were washed to remove excess probes. Purified target-probe complexes were aligned and immobilized onto the nCounter cartridge, and the transcripts were counted *via* detection of the fluorescent barcodes within the reporter probe. Raw gene expression data were analyzed using NanoString nSolver analysis software, version 4.0 Background subtraction was performed using the included negative controls included using the default threshold settings. Data normalization was performed using 6 house-keeping genes (*Gapdh, GusB, Hprt, Oaz1, Polr2 and Sdha*), these genes had been selected based on stability of expression levels from previous transcriptomic experiments. Ratios of transcript count data were generated for primary infected epithelial cells (day 2) versus IAV-rechallenged epithelial cells (day 30+2). Genes with an FDR value <0.05 were considered statistically significant.

### RNA sequencing

Samples were pooled from 6-8 mice in each group. Cells were sorted into 50% FCS/RMPI, centrifuged and washed x1 with PBS and 350μl of RLT added to cell pellets. Samples were centrifuged through a Qiashredder and stored at −80C. RNA was extracted using mini or micro-RNA kits as per manufacturer’s instruction. RNA was purified from total RNA (100ng) using poly-T oligo-attached magnetic beads (Life Technologies, CA, USA). Sequencing libraries were generated using NEB Next Ultra Directional RNA Library Prep Kit for Illumina (NEB) in accordance with the manufacturer’s recommendations. Products were purified and quantified using the Agilent high sensitivity DNA assay on the Agilent Bioanalyzer 2100 system. After the clustering of the index-coded samples, each library preparation was sequenced on an Illumina NextSeq™ 500 platform with 2×75bp paired end sequencing. An average sequencing depth of 20M reads per sample was achieved. Reads were aligned to the mouse genome, mm10, with STAR version 2.4.2a (/doi.org/10.1093/bioinformatics/bts635) in two-pass mode, and on the second pass reads mapping to genes were quantified. Both the genomic DNA sequence (Mus_musculus. GRCm38 primary assembly) and gene annotations (GTF file GRCm38.86) were downloaded from Ensembl (Zerbino *et al*. doi:10.1093/nar/gkx1098). Quality control of the samples was carried out with FASTQC (bioinformatics.babraham.ac.uk/projects/fastqc) and FastP used for trimming, Hisat2 for aligning and FeatureCounts to count the reads aligning to the gene regions. Differential expression analysis was performed with DESeq2 and FDR with a p-value of less than 0.05 was considered statistically significant. Further sample quality control, visualization and exploration were performed in R *via* SearchLight2^83^.

### Over Representation Analysis and Network Analysis

For ORA, significantly different genes were compared to a database (STRING11.5) of pre-defined lists of genes. A hyper-geometric test was then used to determine whether each gene-set is enriched or not for the significantly differential genes. We examined the 10 most enriched gene-sets in the significantly upregulated genes. The false discovery rate (FDR) was calculated to correct the p value. The significant ORA terms were defined an FDR < 0.05. For Network Analysis, enrichment analysis was performed using Hypergeometric Gene Set Enrichment on the gene set databases STRING11.5. For each network nodes represent gene sets and edges represent two gene sets with a Szymkiewicz-Simpson coefficient of at least 0.5.

### RNAscope *in situ* hybridization

Naïve wildtype and IAV infected mice were euthanized by cervical dislocation and lungs were removed. Briefly, lung tissues were fixed with a 10% neutral buffered formalin solution (Sigma-Aldrich, UK) for 24-36 hours and subsequently embedded in paraffin. Lung tissue was cut with a microtome (Leica) at 6µm per section and mounted on Superfrost slides (Fisher Scientific). Slides were stored at R/T until use, then baked at 60°C for one hour prior to commencing the RNA scope protocol. Pre-treatment of tissue sections and RNAscope ISH was carried out according to manufacturer protocols (Advanced Cell Diagnostics, Inc) with protease for 30 mins, followed by heat induced target retrieval, using the RNAscope Red 2.5 kit (ACD-Biotechne; Cat# 322350) with Mm-SpiB (408789), DapB (Cat# 310043) and Mm-Ubc (Cat# 310771). For X31-IAV RNAscope, probe 1226621-C1 (ACD) was used, slides were baked at 37°C overnight and pre-treated, as above, combined with boiling for target retrieval.

### Spatial transcriptomics using Nanostring GeoMx Digital Spatial Profiler (DSP)

X31-IAV RNAscope was used to select virus positive and negative areas from FFPE lungs of each mouse and these cores combined to generate a Tissue Microarray. Two serial 5mm thick slices were cut, one slide tested to confirm virus presence/absence by RNAscope and the second slide used for GeoMX DSP. The GeoMx DSP experiment consisted of slide preparation, with the nuclei stained with Syto13, tissue hybridization with UV-photocleavable probes (Mouse Whole Transcriptome Atlas panel of probes corresponding to 19,963 genes, Nanostring), slide scanning, region of interest (ROI) selection, probe collection, library preparation, sequencing, data processing and analysis. Detailed slide preparation has been previously described^84^.

#### Region of interest and area of illumination selection and probe retrieval

Slides were scanned and imaged at ×20 magnification using the GeoMx DSP with the integrated software suite. Virus positive and negative ROIs were selected using the polygon segmentation tool with reference to the RNAscope, 161 ROIs were selected in total. Images were then used to identify multiple ROIs on which the instrument focuses UV light (385 nm) to cleave the UV-sensitive probes with the subsequent release of the hydridized barcodes with the subsequent aspirate collected into a 96-well DSP collection plate.

#### Library preparation, sequencing and data analysis

Libraries were prepared using GeoMx Seq Code primers (NanoString) and 1× PCR Master Mix (NanoString) and AMPure XP purification. Library quality was checked using an Agilent Bioanalyzer. The libraries were run on an Illumina NovaSeq sequencing system (GeneWiz/Azenta). The FASTQ files from sequenced samples were converted into Digital Count Conversion (DCC) files using the GeoMx NGS pipeline on NanoString’s DND platform. The DCC files were uploaded onto the GeoMx DSP analysis suite (NanoString), where they underwent quality control, filtering, and Q3 normalization^84^. Normalised GeoMx data was analysed using R base functions and packages including Searchlight2^83^. ROI of the same type from the same mice were combined for the analysis.

### Immunohistochemistry

Samples were prepared for immunohistochemistry as described for RNAscope. De-paraffinized and rehydrated sections were incubated in 2% hydrogen peroxide/methanol for 15 minutes. Antigen retrieval (pH 7 or 9) was performed using Vector antigen unmasking (low pH) solution (Vector cat #H-3300) and pH 9 (#H-3301) at 100°C for 20 minutes. Sections were blocked with an avidin/biotin kit (SP-2001; Vector Laboratories, Burlingame, CA) for 20 minutes, then with 10% normal goat serum (NGS, Invitrogen) in PBS for 20 minutes. Slides were incubated overnight at 4°C with primary antibodies at 1:200 directed against B220 (Invitrogen 14-0452-82, clone; RA36A3). Slides were washed with PBS and then incubated with biotinylated goat anti-rat IgG (ThermoFisher, 31830) for 1.5 hours. Amplification of antigen was achieved using an R.T.U. Vectastain kit (Vector, PK-7100), and positive cells were visualized by 3,3-diaminobenzidine tetrahydrochloride (Vector, SK-4100). Hematoxylin and Eosin staining was performed as previously described^17^.

### Immunofluorescence

Mice were euthanized by rising concentrations of carbon dioxide (CO_2_) and the vena cava cut. Lungs were perfused with PBS-5 mM ethylenediaminetetraacetic acid to remove red blood cells and 1% paraformaldehyde was used to fix the lungs. Lung inflation was achieved using 1–3 mL 1% warm UtraPure low melt agarose administrated via the trachea. Lungs with solidified agarose were incubated in 1% paraformaldehyde overnight at 4 °C followed by incubation in 30% sucrose for a further 2–5 days at 4 °C. Lung lobes were frozen in Optimal Cutting Temperature (OCT) (Tissue-Tek 4583, UK) and stored at −80 °C. Lungs were sectioned into 10μm slices on a Shandon Cryotome FE (Thermo Scientific 12087159, UK) and mounted onto Super Frost slides (Thermo Scientific). Slides were fixed in 100% cold acetone and stored at −20 °C.

Slides were rehydrated in PBS containing 0.5% bovine serum albumin (BSA) for 5 minutes and incubated with Fc block (24G2) for 30–60 minutes at room temperature. The sections were stained with antibodies at 4 °C overnight: anti-CD4 AlexaFluor647 (BD UK, RM4-5), anti-MHCII eFluor 450 (M5114, ThermoFisher, UK), anti-B220-PE (ThermoFisher, RA3-6B2). Slides were washed in PBS-BSA and mounted with VectorShied (Vector, UK). Images were collected on an LSM880 (Zeiss, Germany) confocal microscope at 20× magnification using ZEN Black (Version 2.3).

### Digital slide scanning

Excised lung tissues for histological analysis were fixed in 10% neutral buffered formalin for 24 hours and paraffin-embedded in a tissue processor (UK, Shandon Pathcentre Tissue Processor). Formalin-fixed and paraffin-embedded tissues were sectioned (6μm) and processed for subsequent staining with H&E using standard protocols. Stained slides were scanned on a Leica Aperio VERSA 8 bright-field slide scanner (Leica Biosystems, UK) at 10X or 20X magnification and were visualized/analyzed using Aperio ImageScope version 12.1.0.5029 (Aperio Technologies Inc., Vista, USA). Inflammatory infiltrate was identified by blinded observer as described previously^17^ and X31-IAV positive areas detected using ISH were measured using Aperio V9 algorithm (Aperio ScanScope XT System; Aperio Technologies).

### *In vitro* stromal infection and T cell interaction assay

CD45 negative cells were isolated from naïve mice or those that had been challenged with IAV-WSN 30 days previously, as per flow cytometry tissue preparation above and CD45+ cells were removed using a CD45-microbeads as per manufacturer’s instructions (Miletny-Biotec). Cells were seeded at 1×10^6^ cells per well of a 48 well plate and allowed to adhere overnight. Approximately 18 hours later cells were infected with IAV-X31 (Multiplicity of Infection (MOI) 1). T cells were isolated from the spleens of mice that had been infected with IAV-X31 9 days previously to generate a pool of IAV-specific T cells using a T cell isolation kit (Stem cell, as per manufacturer’s instructions). T cells were cultured with infected and control CD45 negative cells for 24h. Flow cytometry analysis was performed as described above.

### Luminex assay

The culture supernatants from the *in vitro* infected stromal cells were harvested, centrifuged to remove cellular debris, and stored at −80 °C until assayed by Luminex (R&D). Each experimental condition was carried out in duplicate. Chemokine/cytokine levels in supernatants were analyzed by quantification with the Milliplex MCYTOMAG-70K assay; IL1α, IL1β, IL10, RANTES/ CCL5, MIG/ CXCL9, IP10/CXCL10, IL6, IL15, IL13, IL17A, IL12p70, IL12p40, GRO/CXCL1 and Milliplex MTH17MAG-47K-03 assay; IL33, IL17F,IL22 (EMD Millipore) according to the manufacturer’s instructions.

### Statistical Analysis

Data were analysed using GraphPad Prism version 10.01 for Windows (GraphPad, San Diego, California, USA). Data were tested for normality using the Shapiro-Wilk test with (α=0.05). Data are presented as mean ± SEM or median ± IQR depending on distribution and p values were calculated as indicated the in the figure legends. P values < 0.05 were considered statistically significant.

## CRediT statement

JCW: Writing – review & editing, Writing – original draft, Visualization, Investigation, Project Administration, Funding acquisition, Formal analysis, Data curation, Conceptualization. KEH: Writing – review & editing, Investigation, Formal analysis, Data curation. GEF: Writing – review & editing, Investigation, Formal analysis, Data curation. CH: Writing – review & editing, Investigation, Formal analysis, Conceptualization. JC: Writing – review & editing, Formal analysis, Data curation. JSN: Writing – review & editing, Formal analysis. FM: Writing – review & editing, Formal analysis. MP: Writing – review & editing, Investigation, Formal analysis. TP: Writing – review & editing, Investigation. KM: Writing – review & editing, Investigation. EB: Writing – review & editing, Investigation, Formal analysis. JA: Writing – review & editing, Investigation, Formal analysis. GI: Writing – review & editing, Investigation, Formal analysis. VH: Writing – review & editing, Investigation, Formal analysis. CKD: Writing– review & editing, Investigation, Formal analysis. YD: Writing– review & editing, Investigation, Formal analysis. NBJ: Writing – review & editing. MP: Writing – review & editing, Funding acquisition. MKLM: Writing – review & editing, Writing – original draft, Visualization, Project administration, Investigation, Funding acquisition, Formal analysis, Conceptualization.

## Acknowledgements

We thank the staff within the school of Infection and Immunity Flow Cytometry Facility, Glasgow Imaging Facility, Biological Services and Polyomics as well as the team of the Veterinary Histopathology Service at the University of Glasgow for technical assistance. We acknowledge the laboratory of Professor Alain Townsend at the University of Oxford for providing the SFLU virus and we would like to thank Dr. Tiong Tan for helpful discussion.

## Financial Support

This work was supported by the Wellcome Trust (210703/Z/18/Z, awarded to M.K.L.M, and 226141/Z/22/Z awarded to MP); a Rosetrees’ Trust Seedcorn Grant (Seedcorn2020\100017, awarded to J.C.W), a Medical Research Council (MRC) Harwell GEMM grant, (GEMM8 2317, awarded jointly to J.C.W and M.K.L.M), and MRC (MC_UU_00034/4, awarded to MP) Wellcome Trust Institutional Strategic Support Funds (097821/Z/11/B awarded to C.H and M.K.L.M and 204820/Z/16/Z awarded to J.C.W), and a University of Glasgow PhD scholarship to G.E.F.

## Conflict of interest statement

The authors have declared that no conflict of interest exists.

## Data availability

RNAseq data are available via GEO GSE278132, Nanostring and GeoMx data are included in Extended data excel and csv files, other primary data available on request to the corresponding authors.

---

**Supplementary Figure 1:**
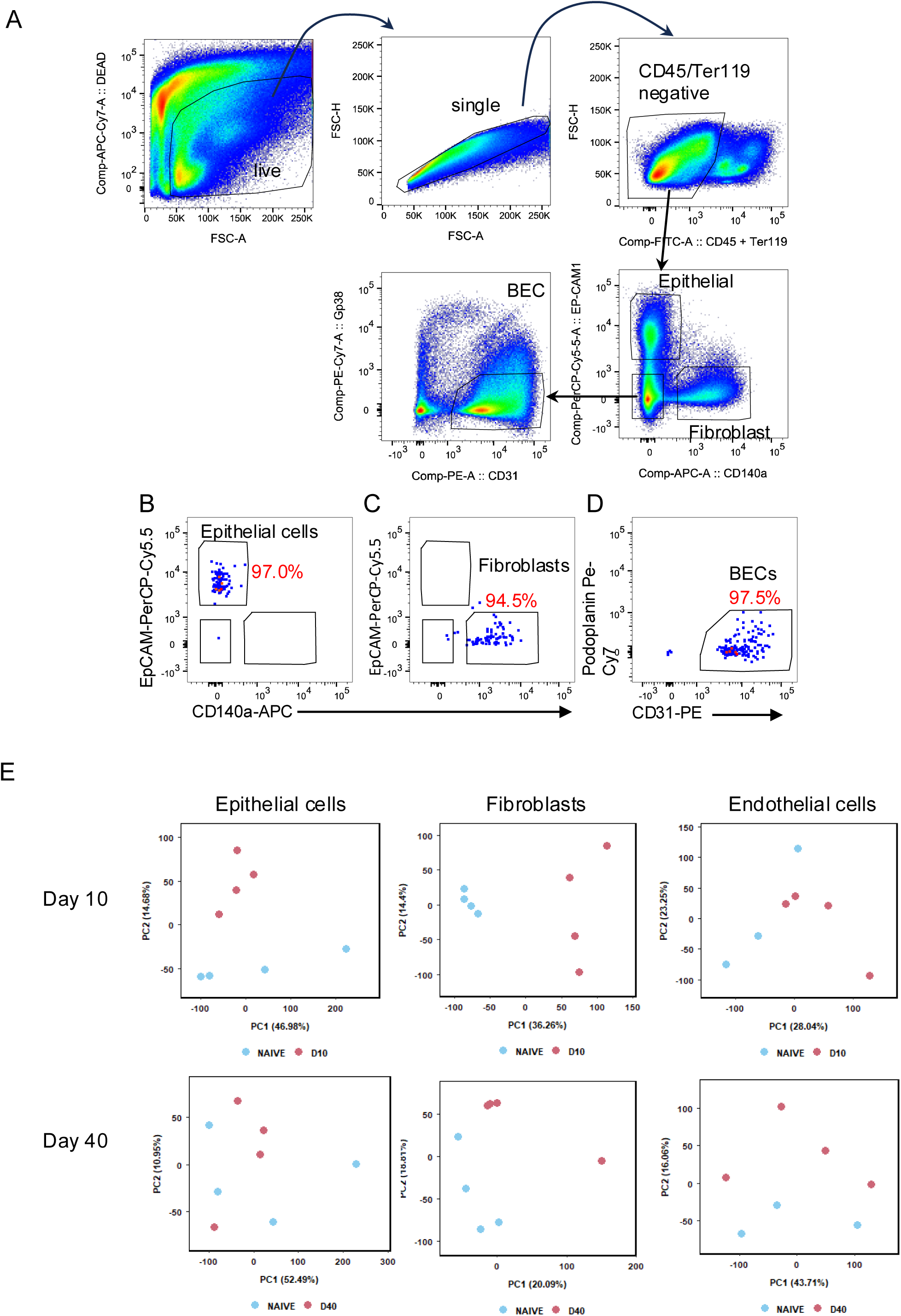
Gating strategy for lung structural cells, sort purity for RNA sequencing, and PCA plots for day 10 and 40 post-infection Sorted stromal gating strategy, sort purity for RNA sequencing and PCA. A: Lung structural cells gated as indicated for FACS sorting. B: Post sort epithelial cells, C: post sort fibroblasts, D: post sort BECs. E: Principal Components Analysis (PCA) for day 10 and day 40 in which each component describes a proportion of the total underlying variation between genes and samples (ordered by the % of the underlying variation that they describe).

**Supplementary Figure 2:**
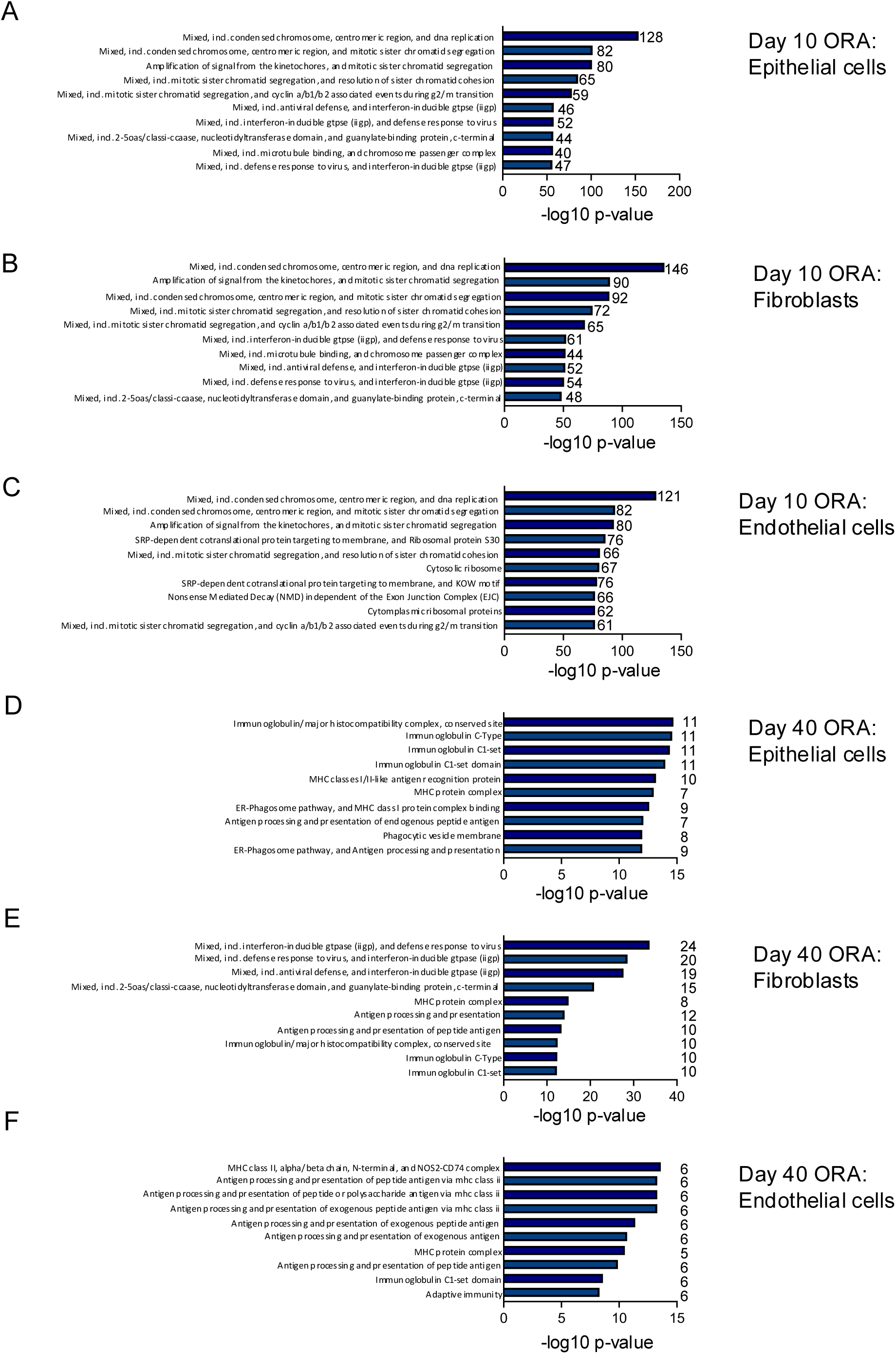
ORA analysis of DEGs in lung structural cells 10 and 40 days following IAV infection. RNA sequencing was performed on sorted lung epithelial cells, fibroblasts, and blood endothelial cells from naïve and C57BL/6 mice infected with IAV 10 (A-C) or 40 (D-F) days previously. Over Representation Analysis (ORA) bar charts showing ten most enriched gene-sets when using significantly upregulated genes with numbers next to each bar showing the number of genes in the pathway differentially expressed. To correct for multisampling a Benjamini-Hochberg correction was applied. Gene sets with an adjusted p-value of 0.05 and an absolute log2fold enrichment above 1 were considered significant.

**Supplementary Figure 3:**
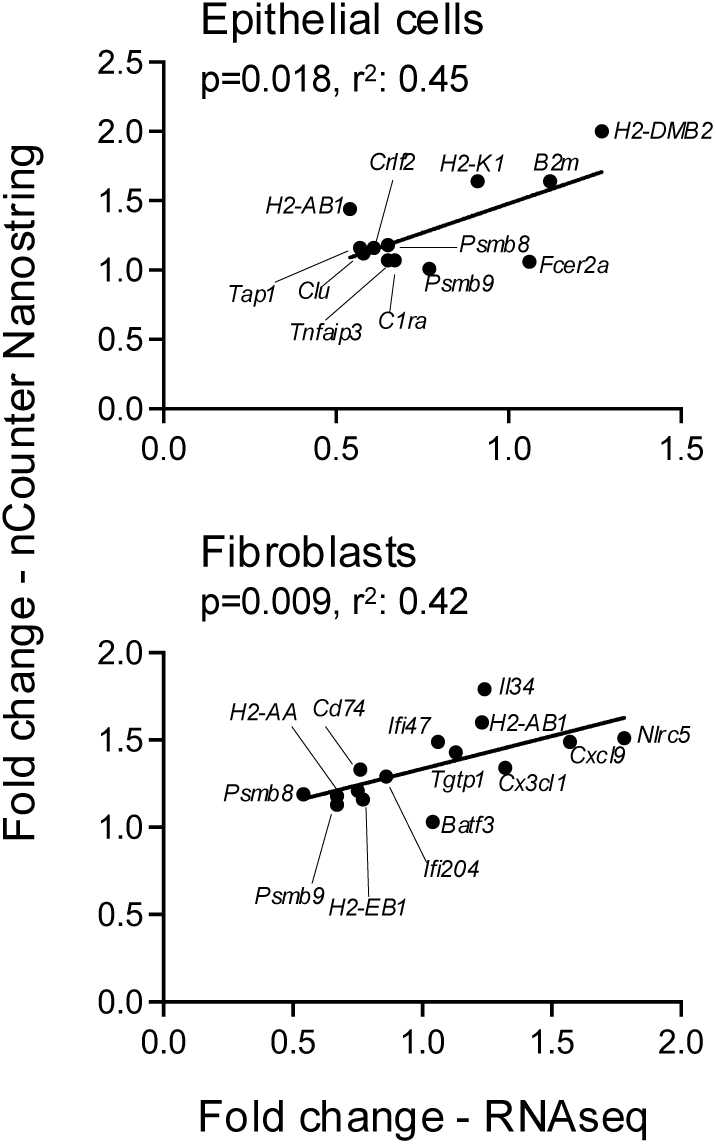
Correlation between RNA-seq and nCounter Analysis of epithelial and fibroblast gene expression following IAV infection. C57BL/6 mice that were either naïve or infected with IAV-WSN and lung epithelial cells and fibroblasts FACS sorted 30 or 40 days later. Gene expression was analyzed either by RNAseq or in a separate experiment by nCounter Nanostring analysis. Selected genes were shared between RNAseq DEG for each cell type and genes in the nCounter Nanostring mouse immunology panel. The data were analyzed using a Pearson correlation and lines on graphs shows the best fit based on linear regression analysis.

**Supplementary Figure 4:**
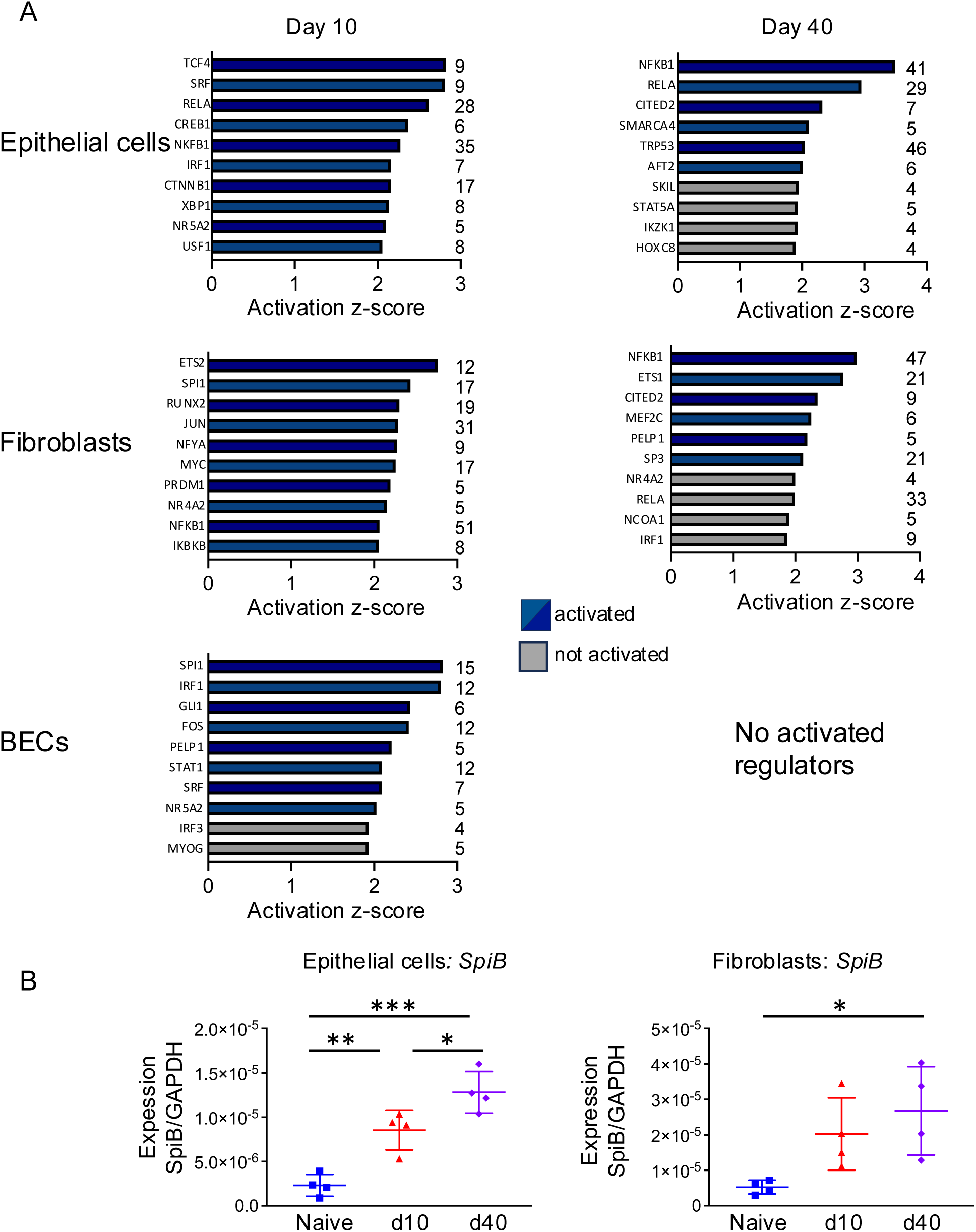
Potential upstream regulators of DEG in lung structural cells post-IAV. A. RNA sequencing was performed on sorted lung epithelial cells, fibroblasts, and blood endothelial cells from naïve and C57BL/6 mice infected with IAV 10 and 40 days previously and potential upstream regulators identified by Activation z-score and the number at the end of each bar shows the number of genes activated in each pathway. B. qPCR data on sorted cells showing absolute copy numbers of SpiB normalized to GAPDH in lung epithelial cells and fibroblasts. *P <0.05 ** P <0.01 and *** P <0.001. Data tested by an ANOVA followed by a Tukey’s multiple comparison test.

**Supplementary Figure 5:**
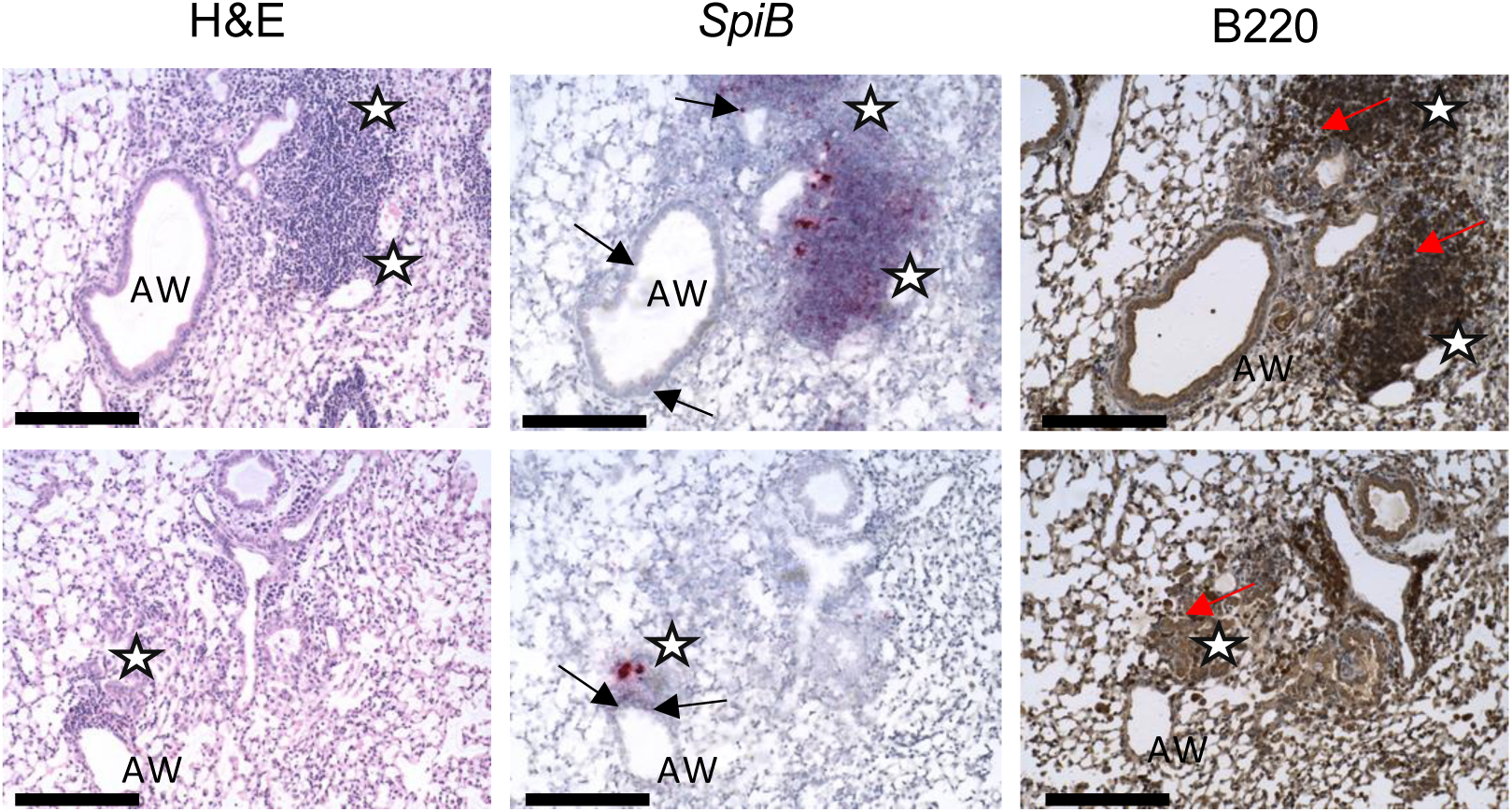
SpiB+ immune cells located in clusters are also B220 positive. Immunohistochemistry and RNA-scope showing SpiB+ airway epithelial cells are located near B220+ immune cell clusters in the IAV infected lung at day 40 post IAV infection. Images taken at 20x magnification, scale bar 200µm. Airways (Aw), SpiB+ epithelial cells (black arrows), inflammatory foci (labelled with stars) and B220+ cells (red arrows).

**Supplementary Figure 6:**
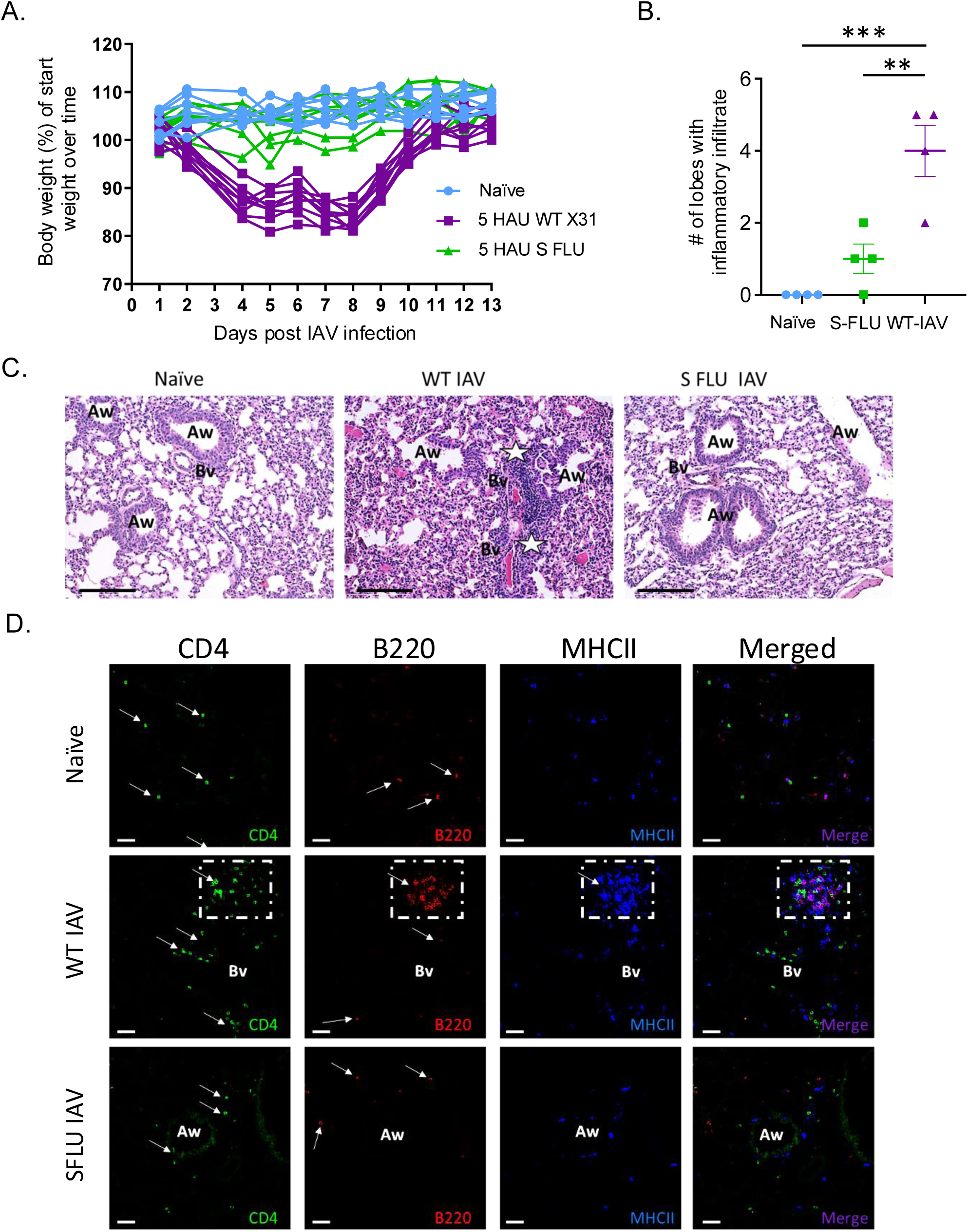
Immune cell clusters do not form following infection with a replication deficient IAV virus (S-FLU) C57BL/6 mice were infected i.n with IAV (X31 strain, either wild type or single cycle (S-FLU)) for 30 days. A: Weight loss graph showing percentage of total body weight over time post infection (n=9) mice per group (naïve, WT-IAV and S-FLU). Data are combined from two independent experiments. B: Histological analysis of inflammation in naïve, WT-IAV and S-FLU infected mice at day 30. Data are normally distributed and shown as mean +/- SEM. One way-ANOVA with Šidák’s multiple comparison, **P <0.01, ***P <0.001. C: H&E staining showing airways (Aw), blood vessels (Bv) inflammatory foci/immune cell clusters (labelled with white stars) in naïve, WT-IAV and S-FLU infected mice at day 30. All images taken at 200x magnification, scale bar 200µm. D: Immunofluorescent staining shows the localization of CD4 T cells (green), B cells (B220+, red) and antigen presenting cells (MHCII, blue) in lung sections from naïve and IAV infected mice culled at day 30 post infection. All images taken at 200x magnification, scale bar 200µm. Airways (Aw), blood vessels (Bv) inflammatory foci/immune cell clusters (labelled with dashed white box). Positive cells are indicated by white arrows.

**Supplementary Figure 7:**
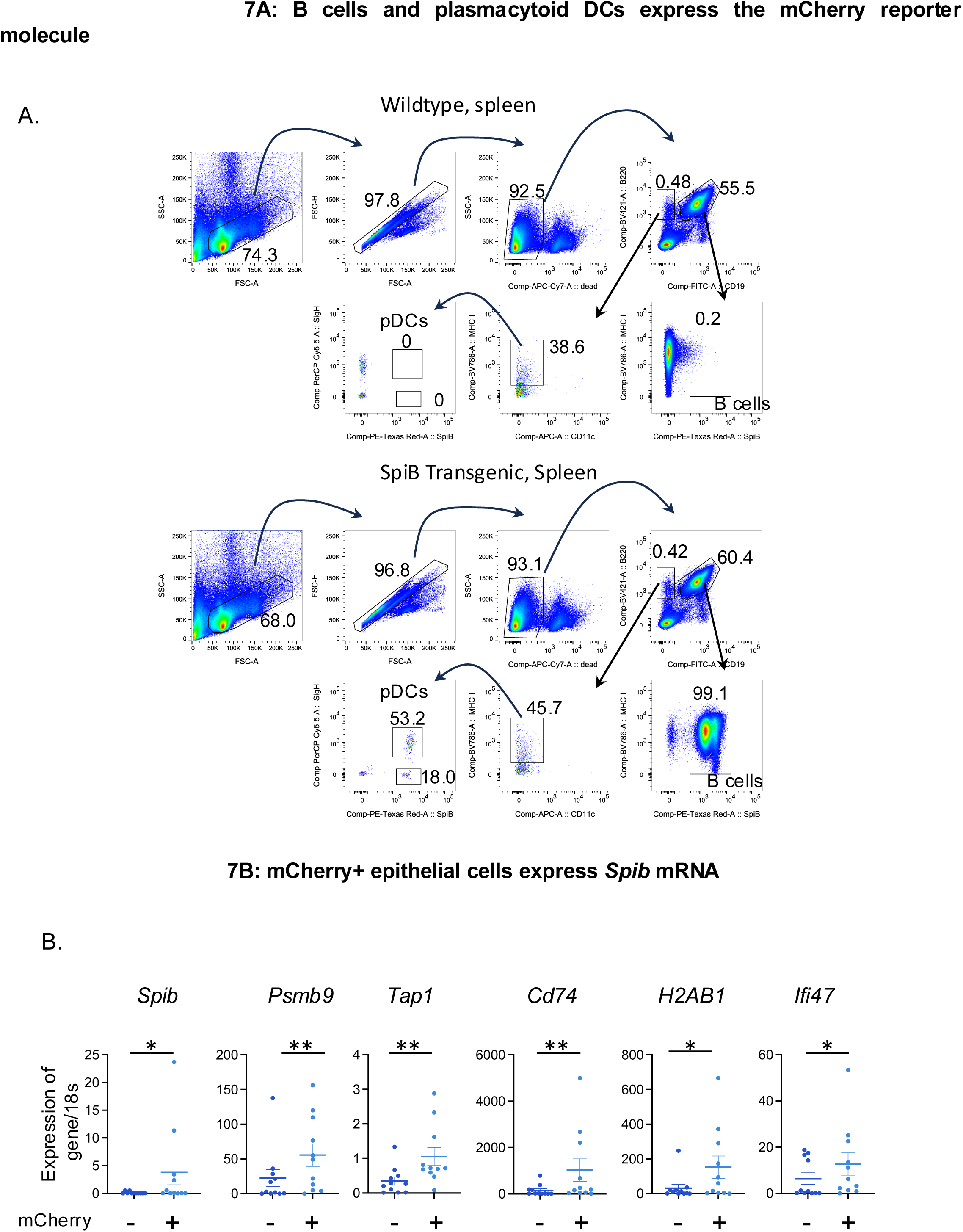
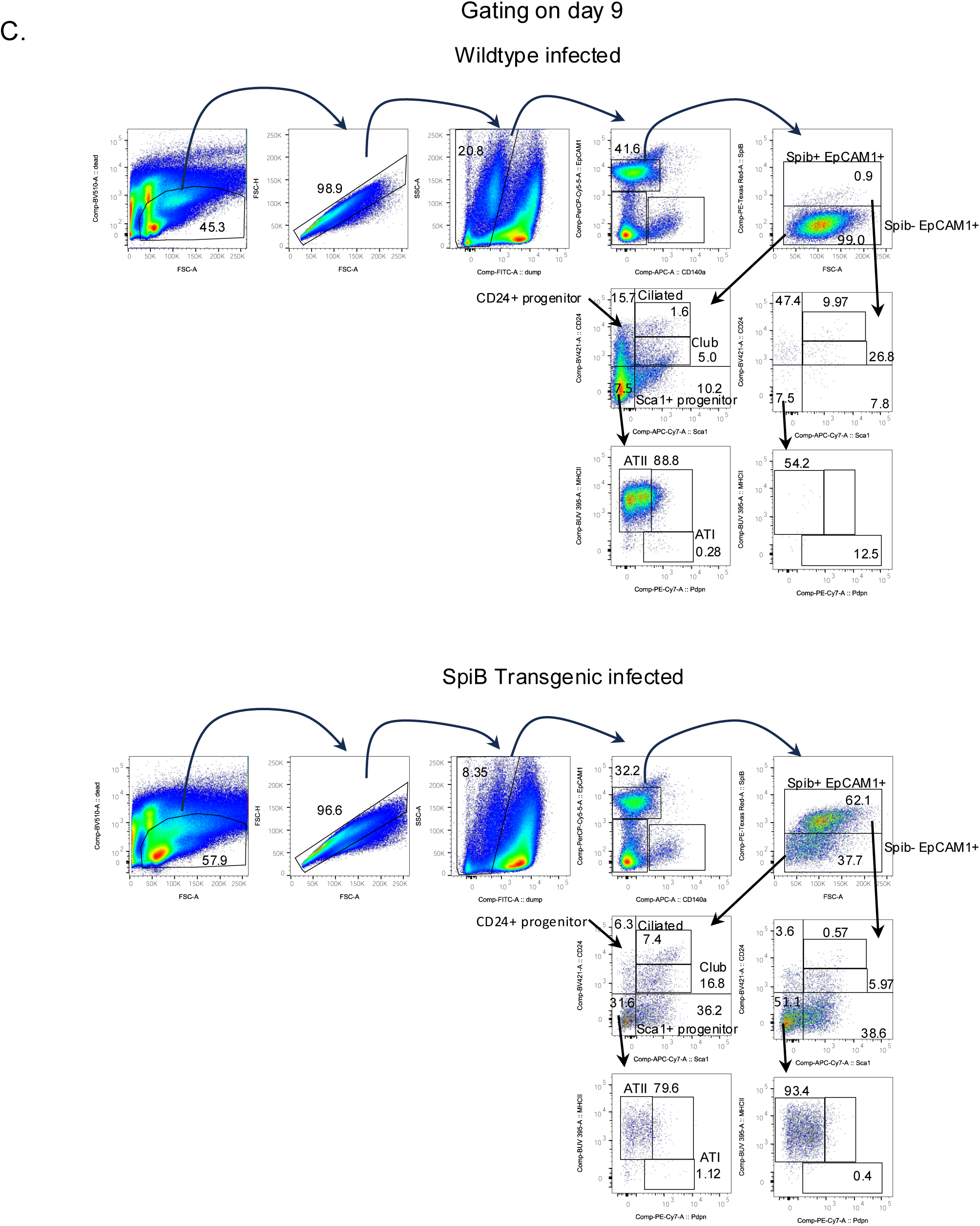
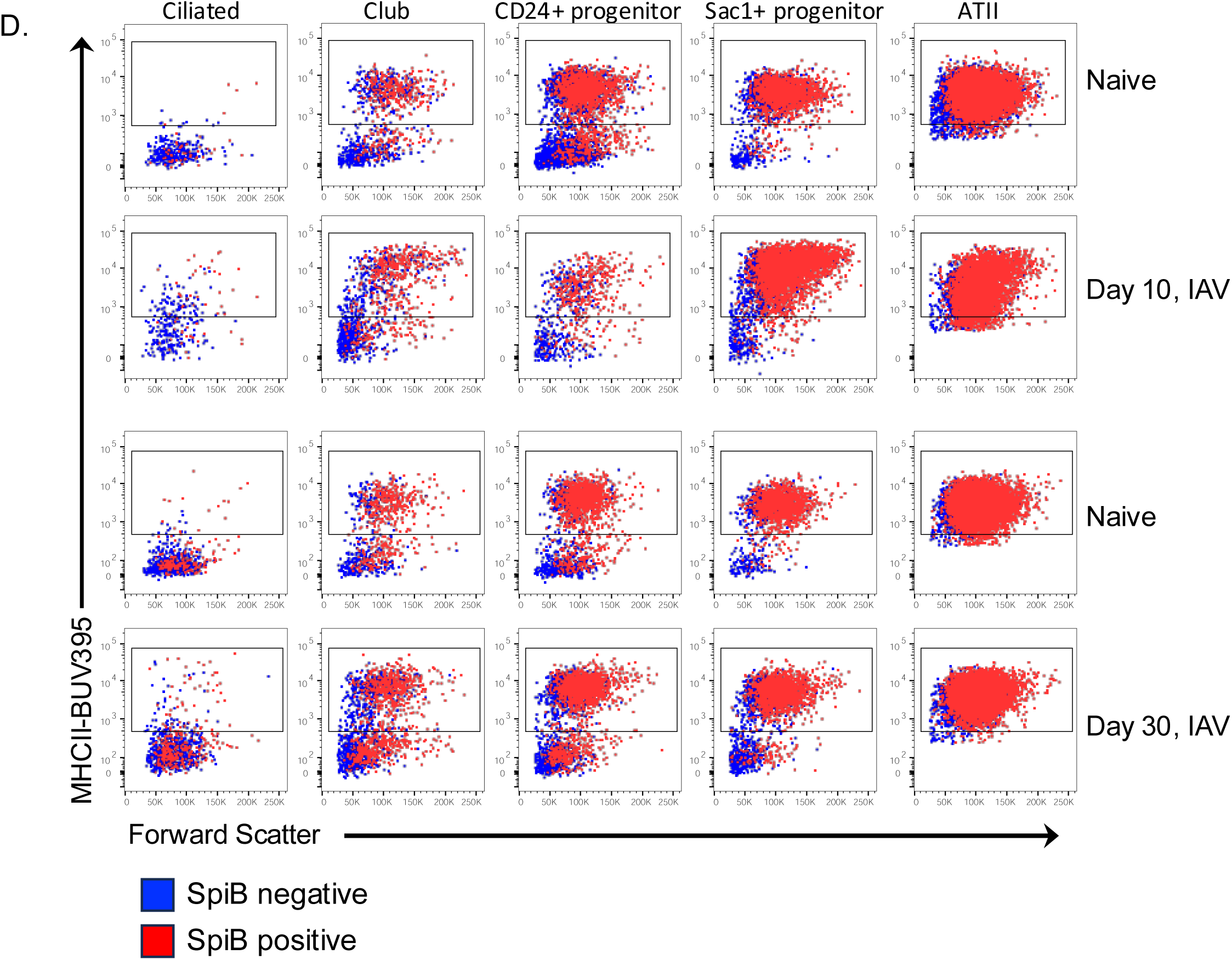
SpiB reporter validation and gating strategy for lung epithelial cell subsets. A: Spleen cells from naïve SpiB-mCherry reporter mice and wildtype C57BL/6 mice were analysed by flow cytometry and B cells and plasmacytoid DCs identified. Numbers indicate percentages within the indicated gates. B: SpiB-mCherry positive and negative cells were FACS sorted based on SpiB+ and SpiB negative EpCAM+ cells based on the initial gating strategy described in C. cDNA was prepared from isolated RNA and examined by semi-quantitative PCR and normalised using the house keeping gene, 18s. Data are not normally distributed and tested via Wilcoxon tests, p<0.05=*, p<0.01= **. C: SpiB-mCherry reporter mice and transgenic negative (Tg neg) littermates were infected i.n. with IAV (WSN) on day 0 and culled at day 9 post infection. Lung cells were gated on live, single, lineage negative (CD45/31, CD140a), EpCAM1+ epithelial cells that were either mCherry (SpiB) positive or mCherry (SpiB) negative. Subsets were then further identified: Ciliated cells (CD24hiSca1hi), club cells (CD24+Sca1+), CD24+ progenitors (CD24+Sca1-) Sca1+ progenitors (Sca1+CD24-), AT1 (CD24-Sca1-PDPN+) and ATII (CD24-Sca1-PDPN-MHCII+). Representative FACS plots are shown for (B) Tg negative littermates and (B) SpiB reporter mice. Numbers indicate the percentage within each gate. D: Representative plots of the SpiB-mCherry negative (blue) and positive (red) epithelial cell populations showing MHCII expression, percentages MHCII positive cells graphed in Figure 4.

**Supplementary Figure 8:**
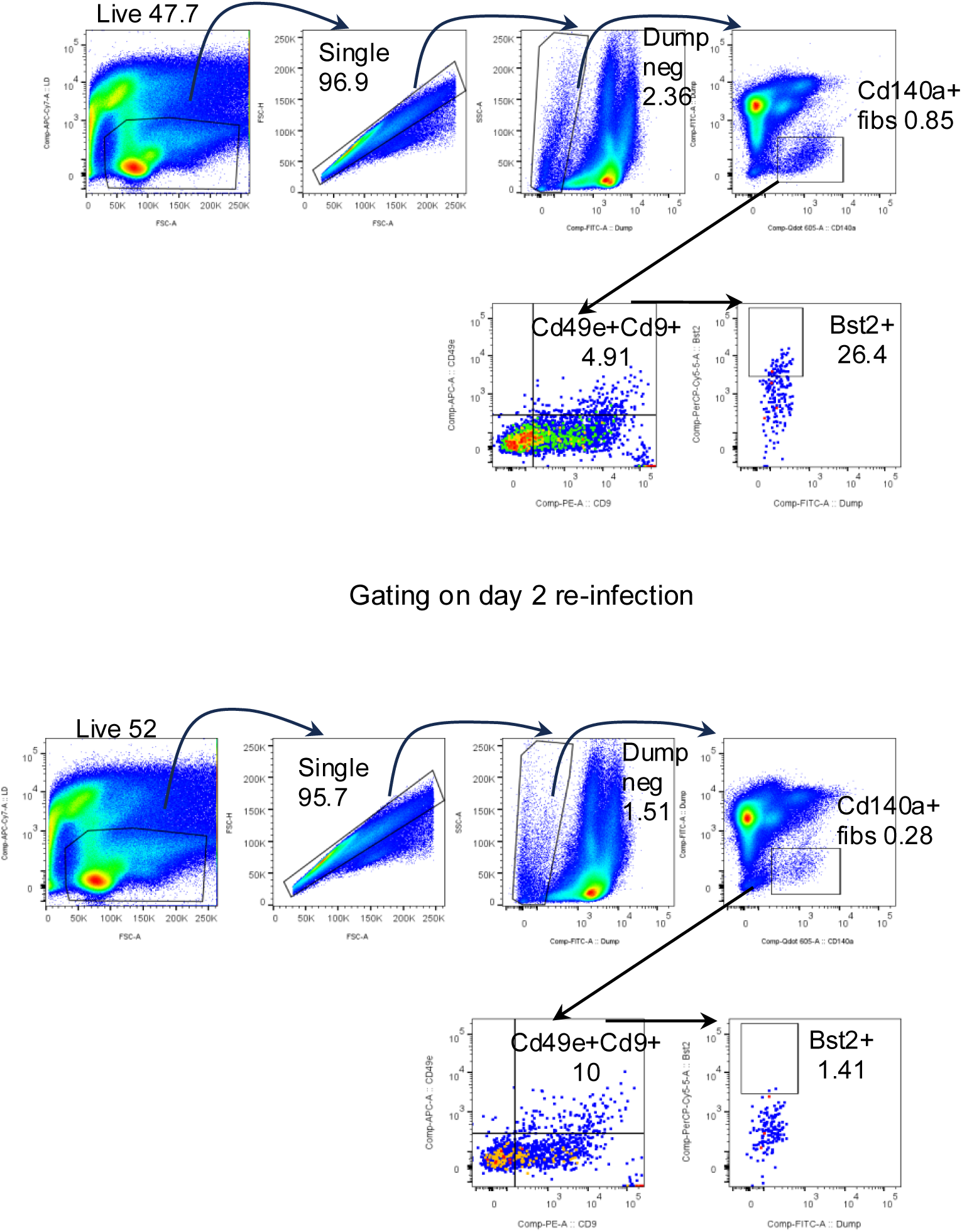
Gating strategy for identification interferon responsive fibroblasts. C57BL/6 mice that were either naïve or infected 30 days earlier with IAV-WSN, were infected with IAV-X31, their lungs harvested 2 days later, and the presence IFN responsive fibroblast populations was examined by flow cytometry. Cells are gated as shown in (A) primary d2 and (B) re-infected d2. Lung cells were gated on live, single, lineage negative (CD45/31/EpCAM negative), CD140a+ fibroblasts that were CD49e+CD9+ and then further gated on Bst2+ versus dump negative populations.

**Supplementary Figure 9:**
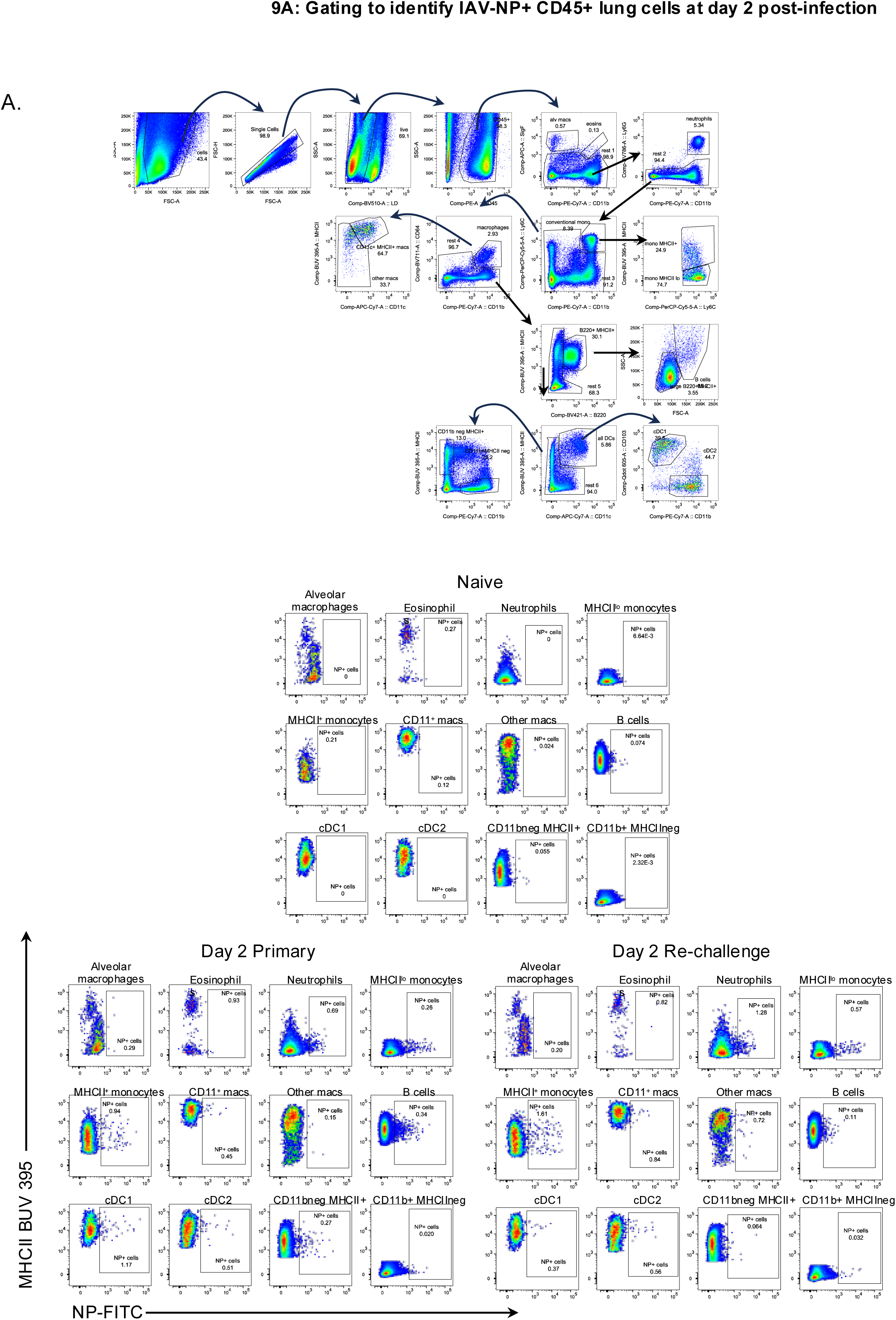
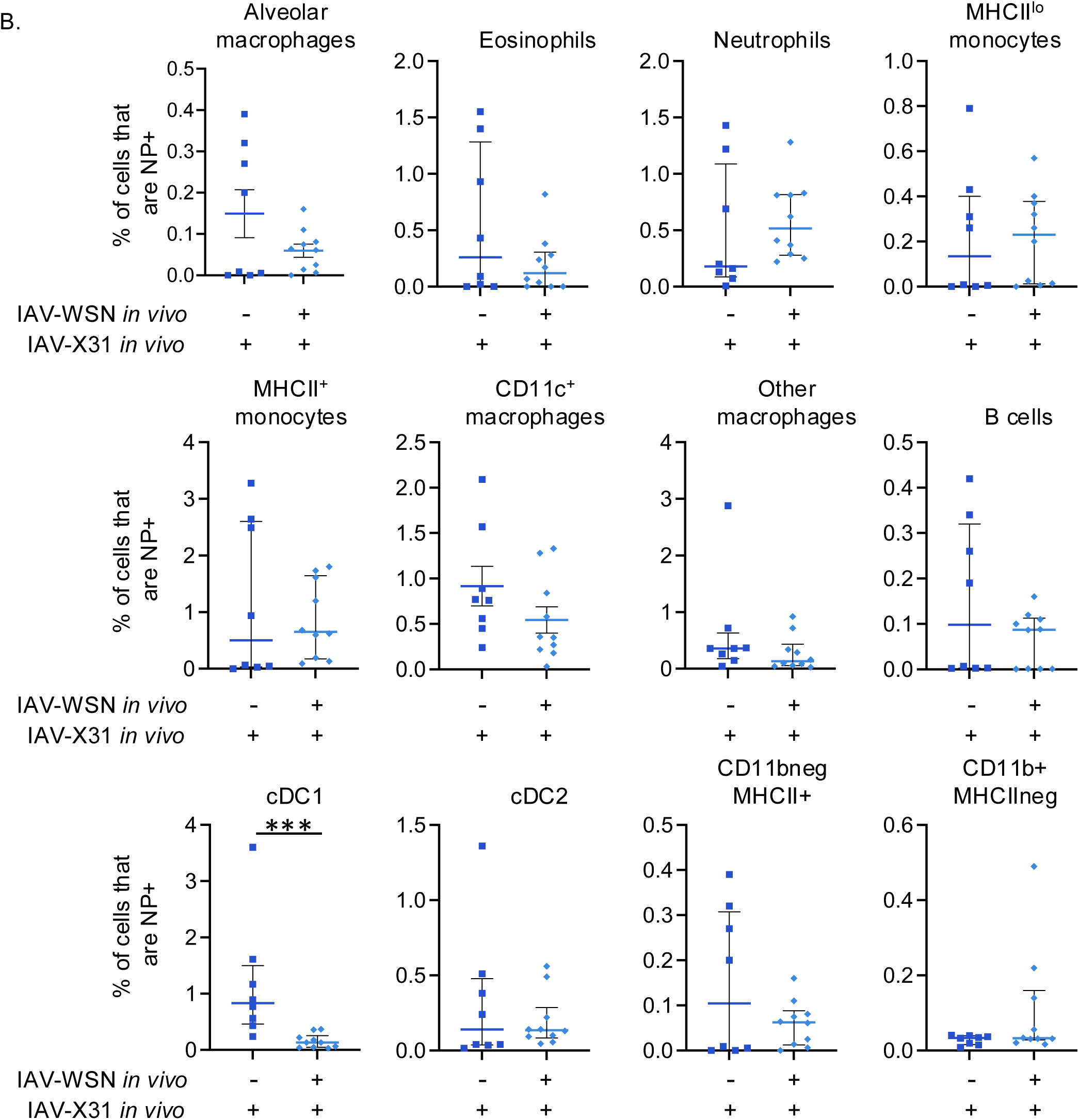
Reduction in viral titers is not due to increased uptake of IAV-NP antigen by APC after IAV re-challenge. C57BL/6 mice that were either naïve or infected 30 days earlier with IAV-WSN, were infected with IAV-X31, their lungs harvested 2 days later, and the presence of IAV-NP within the indicated CD45+ populations examined by flow cytometry. Cells are gated as shown in (A) and combined data shown in (B). Data are from two experiments with a combined number of (n=8) for X31 primary and (n=9) for re-infected groups with each symbol representing a mouse and the horizontal line showing the median for all apart from CD11c+ macrophages where the line shows the mean, error bars are interquartile range expect from CD11c+ macrophages in which they show the SEM. All data apart from CD11c+ macrophage are not normally distributed and differences tested by Mann Whitney, expect from CD11c+ macrophages in which differences tested by a T-test, p<0.001=***.

**Supplementary Figure 10:**
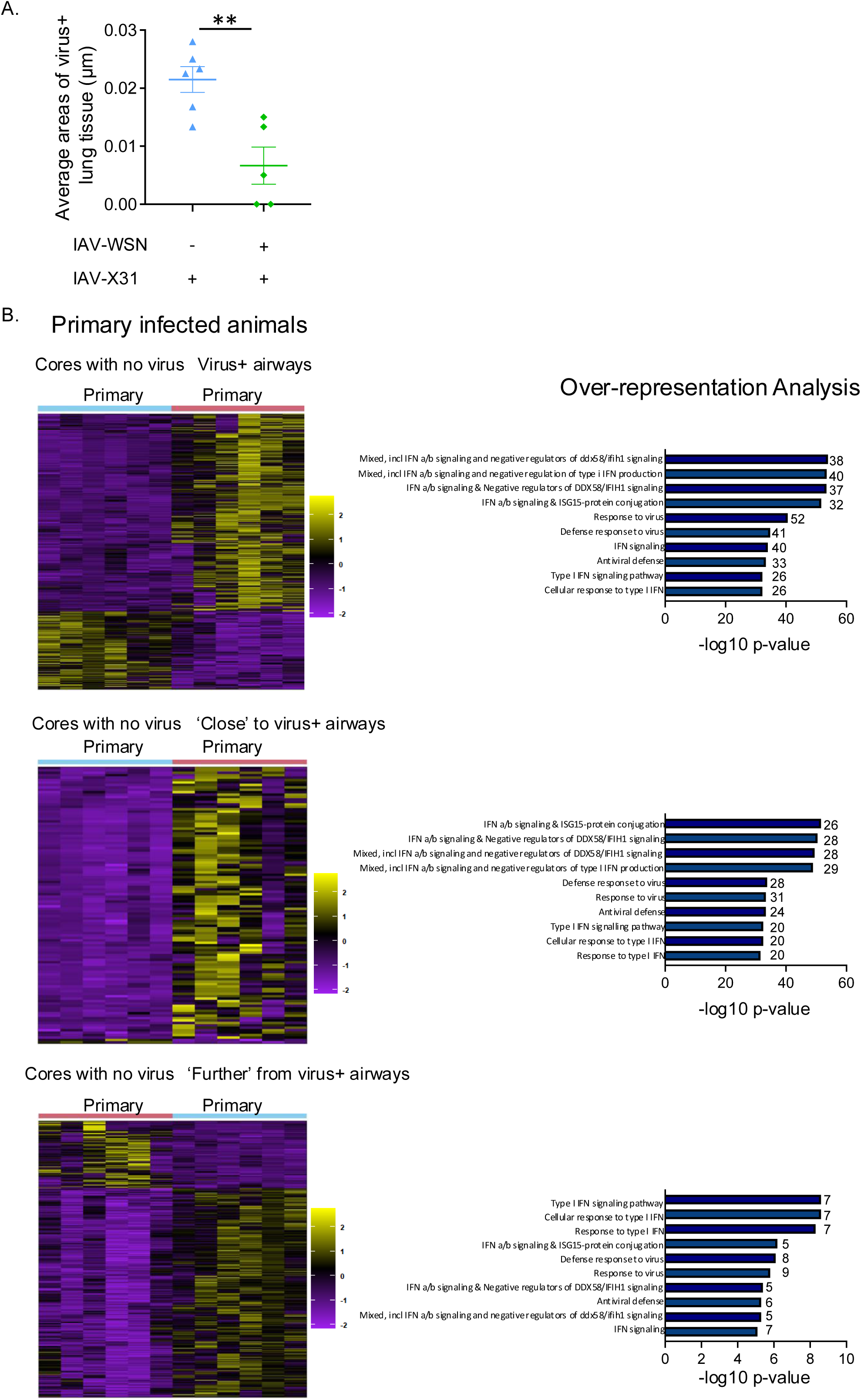
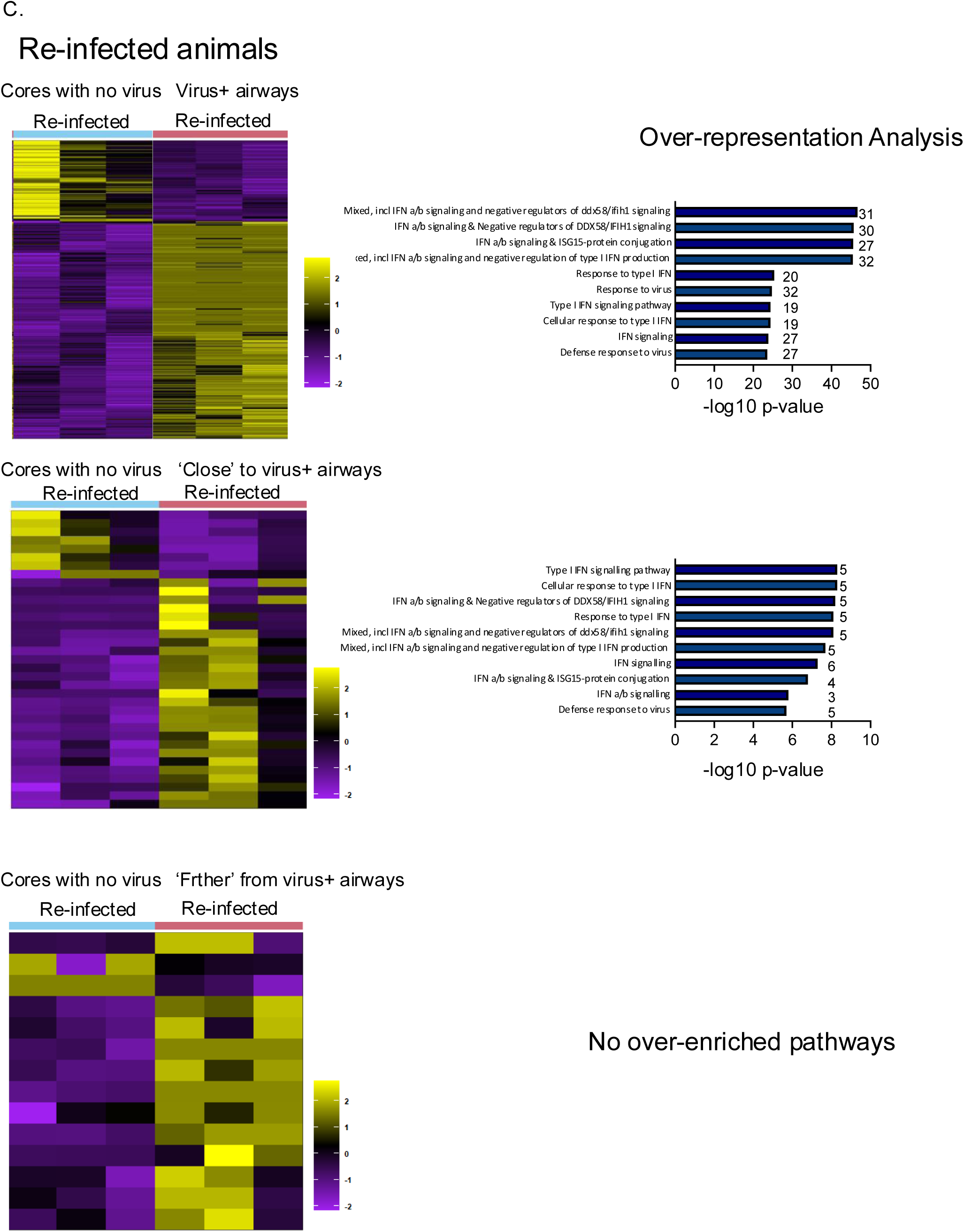
Primary and re-infected mice have a type I IFN response signature in virus+ airways. C57BL/6 mice that were either naïve or infected 30 days earlier with IAV-WSN, were infected with IAV-X31, their lungs harvested 2 days later and embedded in paraffin. Virus positive and negative airways from 6 primary and 3 re-infected mice were identified by RNAscope (A) and analysed by GeoMX. Heatmaps show DEG and enriched pathways in comparisons within primary (B) or re-infected (C) animals between airways from virus negative cores and either: virus+ airways; virus negative areas within the same airway as virus+ cells (Close); or airways adjacent to virus+ airways (Further). Over Representation Analysis (ORA) bar charts showing ten most enriched gene-sets when using significantly upregulated genes with numbers next to each bar showing the number of genes in the pathway differentially expressed.

**Supplementary Figure 11:**
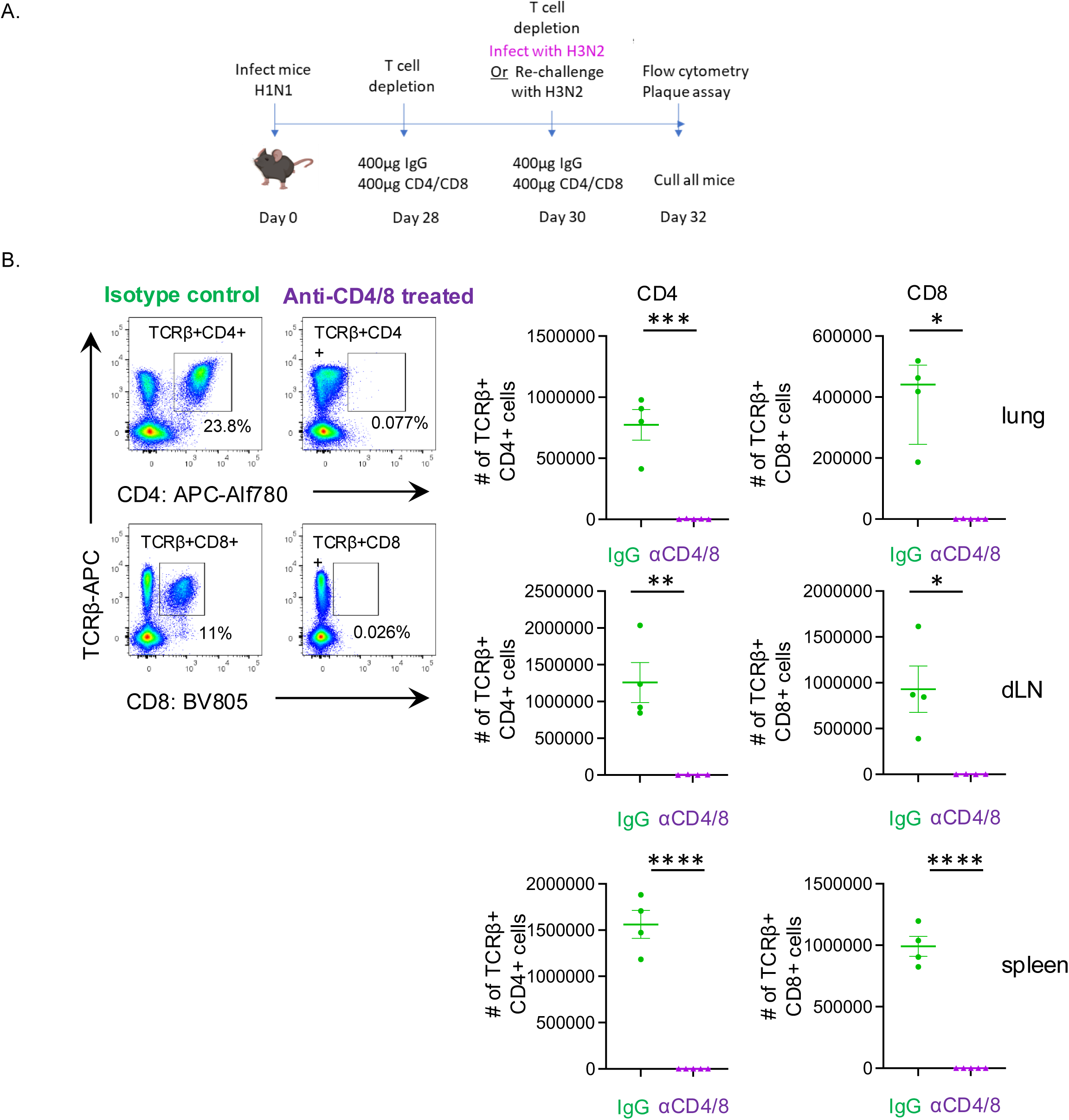
CD4 and CD8 T cells are depleted following two rounds of antibody treatment. (A) Schematic illustrating T cell depletion experimental timeline, C57BL/6 mice were infected with IAV-WSN on day 0 and treated with 400μg isotype control or 200μg each of anti-CD4 (GK1.5) and anti-CD8 (2.43) on days 28 and day 30 when mice were (re)-infected with IAV-X31. (B) Representative FACS plots showing CD4 and CD8 T cell depletion in the lung day 2 post infection, gated on live, single, CD45 iv negative lymphocytes, that are TCRβ+ and numbers of CD4 and CD8 T cells that are TCRβ+ in isotype control (IgG) and T cell deleted groups at day 32. Cell numbers from the lung, draining lymph node and spleen in IgG treated group compared to anti-CD4 and CD8 treated groups and compared by Student’s t test apart from CD8 T cells in lung in which data are not normally distributed, tested Mann Whitney. In graphs, each symbol represents a mouse, the horizontal line shows the mean and error bars are SEM, apart from CD8 T cells in lung which shows median and interquartile range. *:p<0.05, **: p <0.01,***: p<0.001, ****: p<0.0001.

**Supplementary Figure 12:**
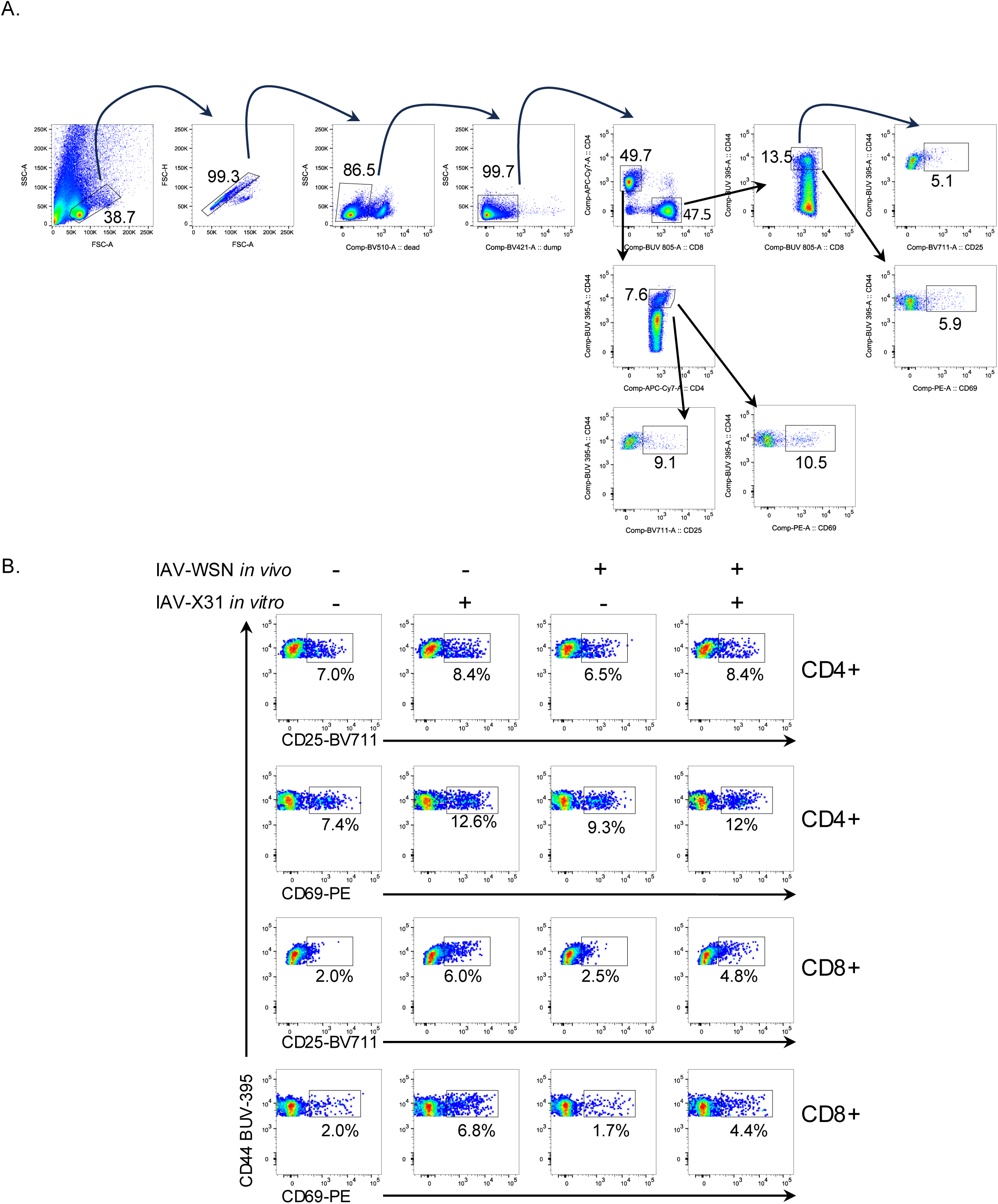
Lung structural cells can present IAV antigens to CD4 and CD8 T cells. Lung CD45 negative cells were isolated from naïve or C57BL/6 mice infected 30 days earlier with IAV-WSN. After 24 hours, the cells were infected with IAV-X31 and co-cultured with T cells isolated from the spleens of mice infected with IAV-X31 9 days earlier. Activated CD4 and CD8 T cells were gated as shown in A and representative FACS plots from each group are shown. Data are representative of two experiments with 4 mice within each naïve and Day 30 IAV group per experiment.

## Notes

### Competing Interest Statement

The authors have declared no competing interest.

### Summary of Updates

One of the extended data sets was incorrect. Revision contains correct Extended data set 4.

